# Saracatinib is an efficacious clinical candidate for fibrodysplasia ossificans progressiva

**DOI:** 10.1101/2020.10.29.360370

**Authors:** Eleanor Williams, Jana Bagarova, Georgina Kerr, Dong-Dong Xia, Elsie S. Place, Devaveena Dey, Yue Shen, Geoffrey A. Bocobo, Agustin H. Mohedas, Xiuli Huang, Philip E. Sanderson, Arthur Lee, Wei Zheng, Aris N. Economides, James C. Smith, Paul B. Yu, Alex N. Bullock

## Abstract

Currently, no effective therapies exist for fibrodysplasia ossificans progressiva (FOP), a rare congenital syndrome in which heterotopic bone is formed in soft tissues due to dysregulated activity of the bone morphogenetic protein (BMP) receptor kinase ALK2/*ACVR1*. From a screen of known biologically active compounds, we identified saracatinib as a potent ALK2 kinase inhibitor. In enzymatic and cell-based assays, saracatinib preferentially inhibited ALK2 compared with other receptors of the BMP/TGFβ signaling pathway, and induced dorsalization in zebrafish embryos consistent with BMP antagonism. We further tested the efficacy of saracatinib using an inducible *ACVR1^Q207D^* transgenic mouse line, which provides a model of heterotopic ossification, as well as an inducible *ACVR1^R206H^* knock-in, which serves as a genetically and physiologically faithful model of FOP. In both models, saracatinib was well tolerated and potently inhibited the development of heterotopic ossification even when administered transiently following soft tissue injury. Together, these data suggest that saracatinib is an efficacious clinical candidate for repositioning in the treatment of FOP, offering an accelerated path to clinical proof of efficacy studies and potentially significant benefits to individuals with this devastating condition.

## Introduction

Current estimates suggest that there are some 7000 inherited rare diseases (defined as affecting fewer than 1 in 2000 people in Europe or 1 in ~1600 in the United States) (1). While their incidence is low individually, rare diseases collectively affect over 7% of the general population. Approximately half of these conditions now have a recognized genetic cause, offering patients hope for new treatments (1, 2). However, the costs and timelines of drug development remain major obstacles for many smaller patient groups, as well as for industry investment. Indeed, fewer than 5% of all rare diseases have an effective pharmacologic treatment, revealing a great unmet medical need (3). One attractive solution for targeting protein kinases is drug repositioning, which enables well-characterized clinical compounds to be tested in novel indications, usually based on clear hypotheses and strong target validation (4–6).

The majority of inherited rare diseases are chronic monogenic conditions that present in early life and cause severely debilitating symptoms as well as reduced life expectancy (7). FOP is a particularly devastating example with significant opportunity for drug repositioning (8). Individuals with FOP become severely disabled due to episodes of heterotopic ossification (HO) that progressively restrict skeletal mobility and reduce lifespan to a median of 56 years (9). Muscle-resident interstitial *Mx1*+ cells mediate intramuscular HO that is dependent upon trauma, whereas tendon-derived *Scx*+ progenitor cells mediate spontaneous and progressive HO in tendons, ligaments and fascia (10). All cases arise from an autosomal dominant germline mutation in the gene *ACVR1* encoding the type I bone morphogenetic protein (BMP) receptor kinase ALK2 (11). Some 97% of patients with the classic FOP phenotype harbour the recurrent gain of function mutation R206H (c.617G>A) (12). This substitution alters the structure of the receptor’s intracellular domain causing aberrant kinase activation and SMAD1/5/8 phosphorylation in response to activin A, as well as hypersensitivity to BMP ligands (13–15).

There are currently no effective treatments for FOP and surgical resection of HO has only proved to exacerbate the condition (8). The discovery that all FOP cases are caused by mutations in ALK2/*ACVR1* has ignited great interest in this pathway as a therapeutic target (11). Prophylactic treatments targeting the ALK2 pathway (13, 16, 17), or associated transcriptional effectors (18, 19), have demonstrated preclinical efficacy in FOP mouse models. The activated kinase domain of ALK2 was first targeted by the small molecules dorsomorphin and LDN-193189 (16, 20). While these tool compounds are not suitable for clinical use, they have demonstrated the potential to inhibit BMP-receptor dependent phosphorylation of SMAD1/5/8 whilst largely sparing the phosphorylation of SMAD2/3 by the activin (ALK4/*ACVR1B* and ALK7/*ACVR1C*) and TGFβ (ALK5/*TGFBR1*) receptors (20). A number of derivatives have also shown selectivity for ALK2 over the other type I BMP receptors ALK1/*ACVRL1*, ALK3/*BMPR1A* and ALK6/*BMPR1B* (17, 21, 22), but considerable time and resource would be needed to optimise these molecules for clinical use.

The protein kinase family provides one of the most successful group of targets for drug discovery. To date, more than 70 small molecule kinase inhibitors have been approved by the FDA and more than 150 are in clinical trials (23, 24). We reasoned that some of these molecules may harbour unreported activity against ALK2, and our unbiased screen proved to reveal a significant number of clinically-tested compounds that bound to ALK2. By far the most potent was the dual SRC/ABL inhibitor saracatinib (AZD0530), which demonstrated low nanomolar inhibitory activity against ALK2. In cells, this compound was also 30-fold selective for inhibition of BMP6 over TGFβ signalling pathways. Most importantly, saracatinib blocked HO in two relevant mouse models. Thus, saracatinib is a promising investigational drug to test in clinical trials as a frontline therapy for FOP.

## Results

### Saracatinib is a potent inhibitor of the protein kinase ALK2

To identify potential drug candidates for application in FOP we used the kinase domain of the BMP type I receptor ALK2 to screen a library of clinically tested small-molecule inhibitors via differential scanning fluorimetry (DSF). Ligands in this assay increase a protein’s melting temperature (*T*_m_ shift) by an amount proportional to their binding affinity (25). The most potent hit was the dual SRC/ABL inhibitor saracatinib (AZD0530, Figure 1A), which induced a large *T*_m_ shift of 13.9°C comparable to that of the leading tool compound LDN-193189 (26, 27) (Figure 1B and Supplemental Table 1). By contrast, the next hit, ASP3026, produced a *T*_m_ shift of just 10.7°C equivalent to the weaker tool compound dorsomorphin (20, 26). As a control for inhibitor selectivity, a counter screen was performed using the kinase domain of TGFβ type I receptor ALK5. Importantly, saracatinib induced a significantly lower *T*_m_ shift with ALK5 of 10.2°C, suggesting that this inhibitor was both potent and selective for BMP vs. TGFβ signalling (Figure 1B and Supplemental Table 1). We also tested saracatinib against ALK2 mutants implicated in FOP disease, finding that all mutants displayed *T*_m_ shifts above 14.5°C, confirming their strong interaction (Table 1). Interestingly, the mutant proteins were all destabilized in their apo states in solution, consistent with structural models showing that FOP-causing mutations break interactions that normally stabilize the receptor’s inactive conformation (14).

**Figure 1.**
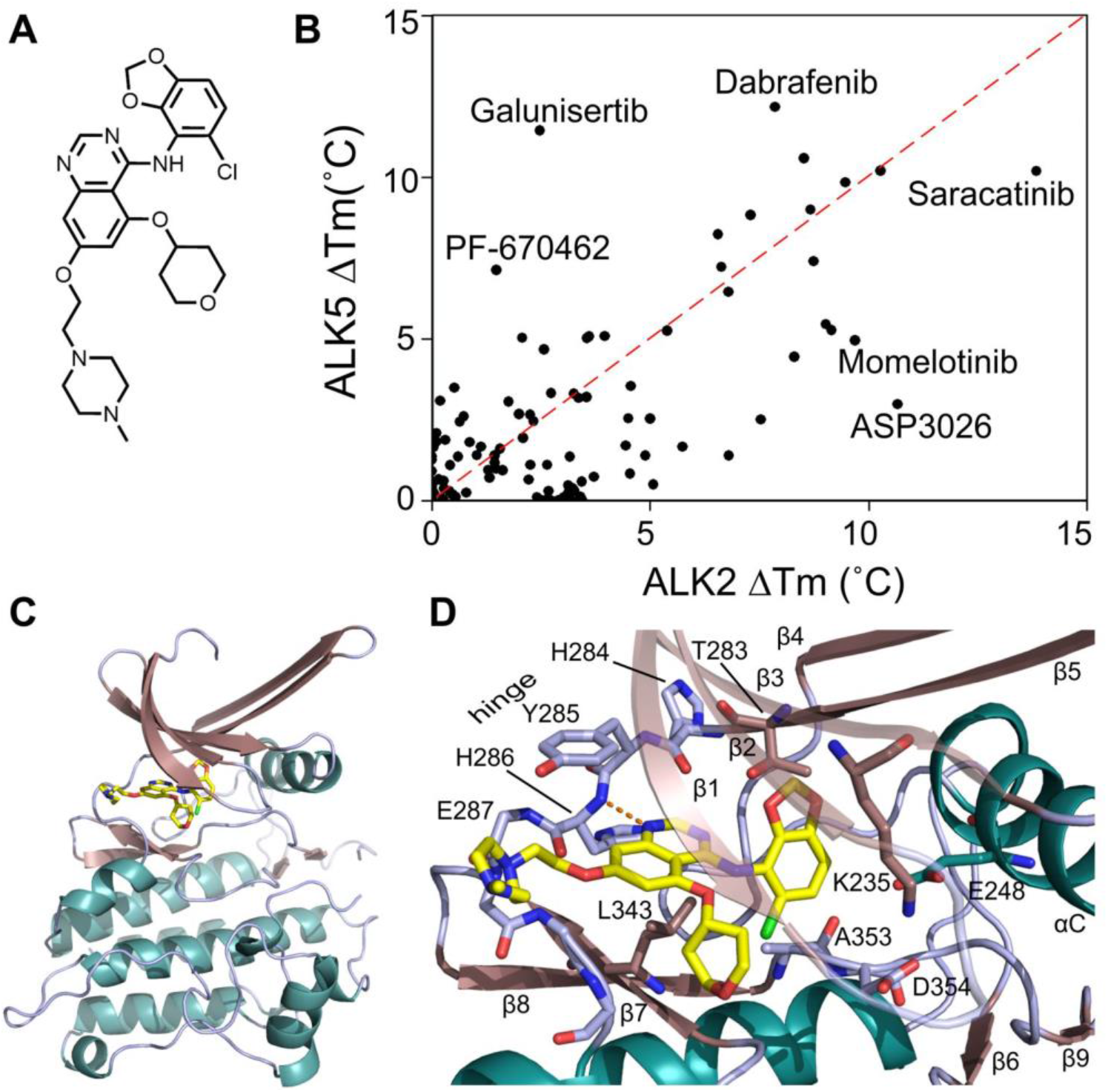
DSF screening identifies saracatinib as a potent inhibitor of ALK2. (**A**) Chemical structure of saracatinib. (**B**) Plot comparing the thermal shift (Δ*T*_m_) of ALK2 and ALK5 in response to different clinical compounds. Selected compounds of interest are labelled including saracatinib. A red dashed line is drawn as an approximate guide to mark equipotency for ALK2 and ALK5. (**C**) Overview of the ALK2 co-crystal structure with saracatinib. (**D**) Interactions of saracatinib with the ATP-binding pocket of ALK2.

**Table 1.** DSF results for ALK2 variants

| ALK2 protein | $T_m$ (°C) | | $\Delta T_m$ (°C) |
| --- | --- | --- | --- |
|  | Apo | + saracatinib |  |
| WT | 42.4 | 56.4 | 13.9 |
| L196P | 40.0 | 57.6 | 17.6 |
| R206H | 40.9 | 55.4 | 14.5 |
| Q207E | 42.2 | 57.6 | 15.4 |
| R258S | 38.1 | 56.4 | 18.3 |
| G328E | 37.0 | 56.0 | 19.0 |

### Saracatinib binds inactive ALK2 in an αC-in conformation

Although still in phase III trials, saracatinib is based on the quinazoline scaffold, which has yielded numerous approved drugs, including gefitinib and vandetanib (28). Interestingly, none of the other quinazoline-based kinase inhibitors tested showed any significant *T*_m_ shift against ALK2 or ALK5 (*T*_m_ shift of 4.5°C or less), indicating that this was a specific property of saracatinib (Supplemental Figure 1, A-C). To explore the molecular basis of this interaction we solved the crystal structure of this inhibitor in complex with the ALK2 kinase domain and refined the structure at 2.66 Å resolution (PDB accession 6ZGC, Figure 1C, see Supplemental Table 2 for diffraction data collection and refinement statistics). Within the ATP-binding pocket, the bound ligand was well resolved by the electron density map (Supplemental Figure 2). While the quinazoline of saracatinib formed a hydrogen bond to the hinge residue His286, specificity was likely derived from the large chlorobenzodioxole moiety which nicely complemented the size, shape and hydrophobicity of the back pocket (Figure 1D). The tetrahydropyran ring was located in the ribose pocket below the glycine-rich loop (β1-β2), whereas the methylpiperazine moiety was extended largely into solvent (Figure 1D).

Structural differences between the ALK2 and SRC complexes with saracatinib were evident from the different conformations of their catalytic domains. The previously determined structure of the SRC complex displayed an inactive αC-out conformation that widened the ATP pocket and rotated the αC residue Glu310 away to solvent, as well as the glycine-rich loop residue Phe278 (Supplemental Figure 3) (29). By contrast, the ALK2 complex exhibited an inactive conformation in which the αC helix was swung inwards to compress the ATP pocket. As a result, the equivalent αC (Glu248) and glycine-rich loop (Tyr219) residues were packed inside the ATP pocket where they mediated additional interaction with the chlorobenzodioxole and tetrahydropyran groups, respectively (Supplemental Figure 3). Thus, further optimization of this chemotype could yield molecules with increased selectivity for ALK2 over SRC family kinases.

Saracatinib was developed to be relatively selective for SRC/ABL kinases compared to some other multi-targeted kinase inhibitors used in oncology today (29). We extended the kinase inhibition profiling for this compound to a panel of 252 human kinases using the Caliper technology (Nanosyn, Inc). Using 100 nM saracatinib, LCK and ALK2 ranked first and second as the most strongly inhibited kinases, followed by SRC, RIPK2 and ABL (Supplemental Table 3). Importantly, ALK2, RIPK2 and TNIK were the only serine/threonine kinases showing >50% inhibition at this concentration, whereas the others were all tyrosine kinases, including paralogs such as ARG (ABL2) and the SRC-family kinases HCK, FYN and LYN. These observations are consistent with determined IC_50_ values for saracatinib against purified activin receptor-like kinases (IC_50_: ALK2, 6.7 nM > ALK1, 19 nM >> ALK3, 621 nM >> ALK4, 3900 nM > ALK6, 6130 nM and ALK5, 6890 nM, Figure 2A). Thus, saracatinib shows both the inhibitory potency and selectivity desired in a clinical candidate for FOP.

**Figure 2.**
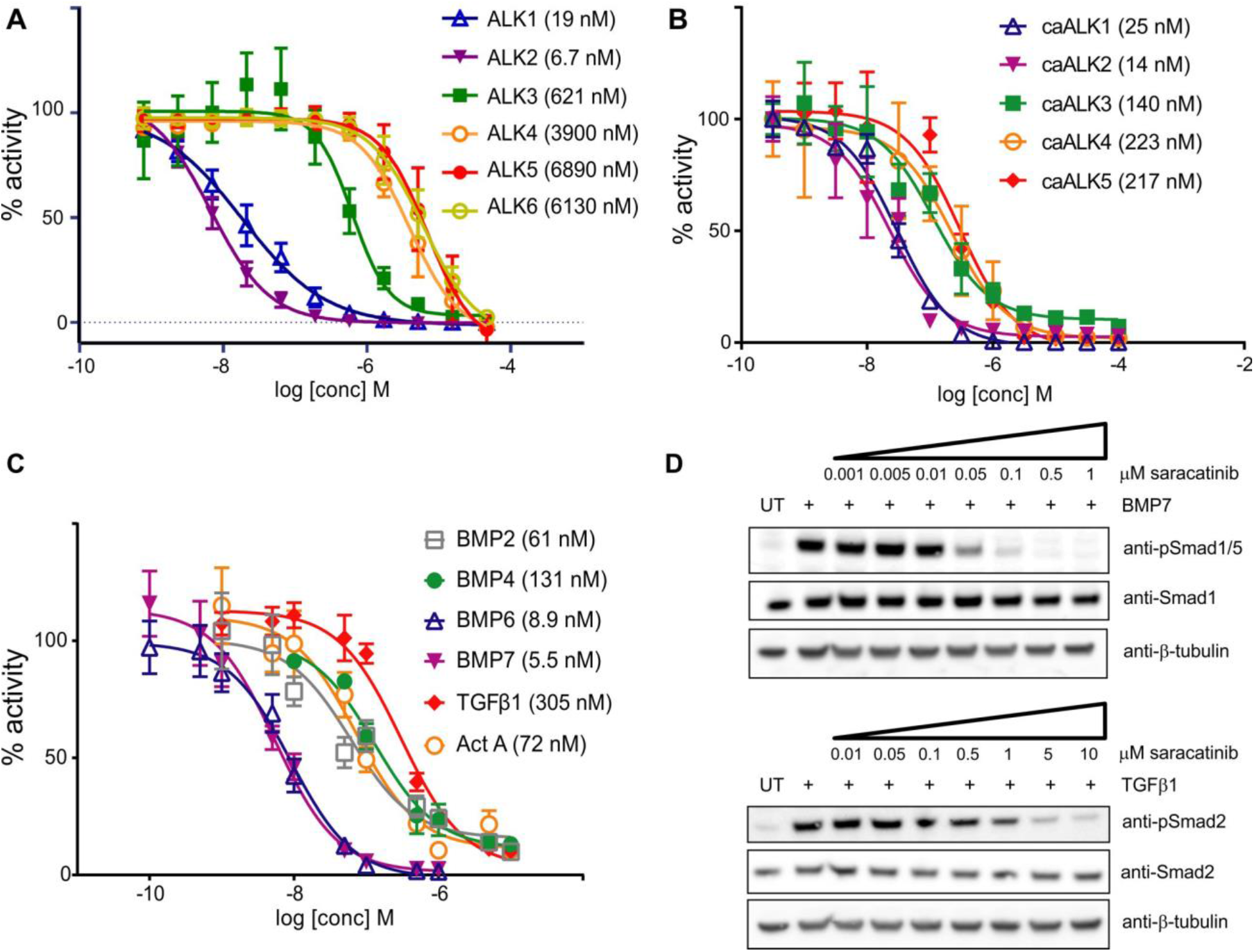
Saracatinib is a selective inhibitor of BMP versus TGFβ type I receptor activity in cells. (**A**) IC_50_ inhibition curves for saracatinib against purified recombinant ALK1-6 kinase domains were determined using a LANCE Ultra TR-FRET kinase assay (PerkinElmer). Reactions contained 10 nM kinase, 50 μM peptide substrate and 10 μM ATP. Data shown are plotted as mean ± S.D. (n=3 independent replicates). (**B**) Representative inhibition curves for saracatinib against constitutively active BMP (caALK1, 2 and 3) and activin/TGFβ (caALK4 and 5) type I receptors, based on the activity of BMP responsive promoter element luciferase (BRE-Luc) and TGFβ responsive luciferase (CAGA-Luc) reporters in C2C12 and 293T cells, respectively. Data shown are representative of more than 3 independent experiments, with data plotted as mean ± S.E.M. (n=3 replicates). (**C**) IC_50_ inhibition curves for saracatinib against the signaling induced by indicated ligands based on the activity of BRE-Luc (BMP ligands) and CAGA-Luc (activin/TGFβ ligands) reporters stably expressed in MDA-MB-231 cells. Data shown are plotted as mean ± S.D. (n≥3 independent replicates). (**D**) Western blot analyses showing the inhibitory activity of saracatinib against BMP7-induced phosphorylation of SMAD1/5, as well as TGFβ-dependent phosphorylation of SMAD2 in C2C12 cells.

### Saracatinib inhibits wild-type and mutant ALK2 in cells

To confirm this selectivity profile in cells, we first measured the effect of saracatinib on gene expression driven by constitutively active (ca-) forms of the receptors. Inhibition of the BMP receptors caALK1, caALK2 and caALK3 was assessed via the activity of a BMP-response element luciferase reporter (BRE-Luc) stably expressed in the C2C12 cell line, whereas the activin (caALK4) and TGFβ (caALK5) receptors were profiled via a CAGA-Luc reporter construct stably expressed in HEK293 cells (Figure 2B). As expected for an inhibitor in clinical use, saracatinib was well tolerated in both cell lines up to a concentration of approximately 40 μM (Supplemental Figure 4). Saracatinib most potently inhibited the BMP receptors caALK2 and caALK1, with IC_50_ values of 14 nM and 25 nM, respectively. Inhibition of BMP receptor caALK3 was more modest (IC_50_ = 140 nM), while inhibition of the activin/TGFβ receptors caALK4 and caALK5 was weaker (IC_50_ ~ 220 nM) (Figure 2B).

For subsequent investigations of ligand-dependent receptor signalling we utilized MDA-MB-231 cells stably expressing either the BRE-Luc or CAGA-Luc construct. Again, the inhibition profile of saracatinib showed significant selectivity towards ALK2 and its preferred ligands BMP6 and BMP7. Saracatinib potently inhibited signalling downstream of BMP6 and BMP7 with IC_50_ values of 8.9 nM and 5.5 nM, respectively, while it was less potent against signalling downstream of BMP2 (IC_50_ = 61 nM) and BMP4 (IC_50_ = 131 nM) as shown in Figure 2C. Canonical activin A and TGFβ signalling were inhibited to a lesser extent (IC_50_ = 72 nM and 305 nM, respectively, Figure 2C). Thus, there was a 30-fold increase in the concentration of saracatinib required to inhibit TGFβ signalling compared to that of preferred ALK2 ligands. A similar pattern was observed in C2C12 cells using specific antibodies to detect receptor-mediated phosphorylation of the substrate SMAD molecules. Western blot analyses revealed that BMP7-induced phosphorylation of SMAD1/5 was completely inhibited by 100 nM saracatinib, whereas the TGFβ-dependent phosphorylation of SMAD2 was only blocked at an inhibitor concentration of 5 μM (Figure 2D). SMAD phosphorylation induced by other BMP and activin ligands was inhibited at intermediate saracatinib concentrations consistent with the luciferase reporter assays (Supplemental Figure 5).

FOP-causing mutations in *ACVR1* induce neo-function in transducing activin A via BMP-receptor associated SMAD1/5/8 (13, 15), and this gain of function appears to be the major pathogenetic mechanism for the formation of heterotopic bone. To test whether saracatinib could also inhibit activin A-induced activation of SMAD1/5, we used primary dermatofibroblast cells derived from FOP patients with the classic *ACVR1^R206H^* mutation or wild-type control cells. Western blot analysis confirmed that phosphorylation of SMAD1/5 in response to activin A was observed only in the FOP patient-derived cells and not in wild type (Figure 3). In the presence of 100 nM saracatinib, this phosphorylation was inhibited with similar efficacy to that shown using the control ligand BMP6 (Figure 3A). An IC_50_ of 15 nM was determined using an in-cell immunofluorescent assay (Supplemental Figure 6), further confirming the ability of this molecule to block the neo-function of ALK2 implicated in the development of FOP.

**Figure 3.**
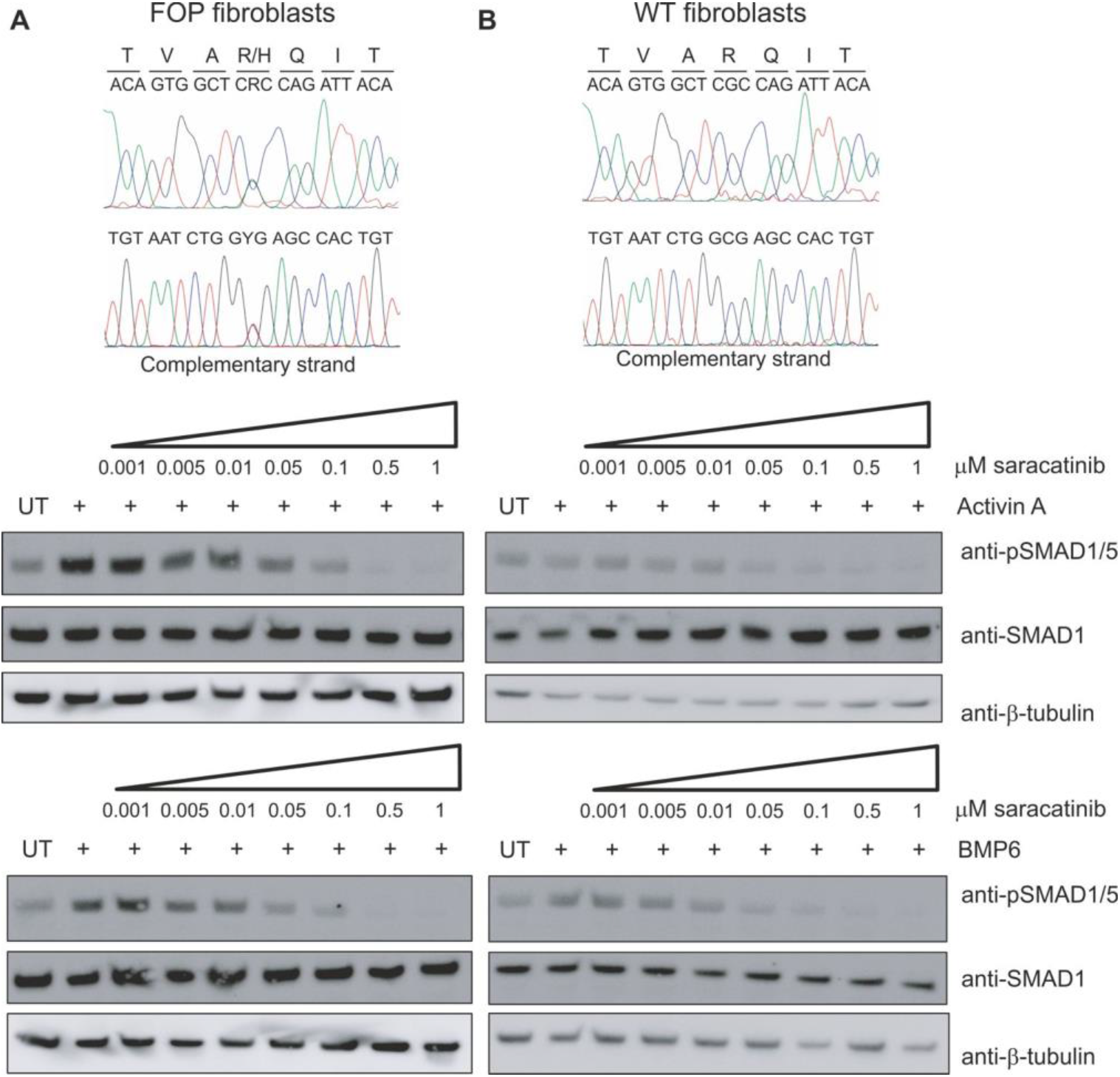
Saracatinib inhibits the neofunction of ALK2^R206H^. Western blot analysis of phospho-SMAD1/5 levels following treatment with saracatinib and either BMP6 or activin A in (**A**) FOP patient-derived fibroblasts cells (GM00513) or (**B**) wild-type fibroblasts cells (ND34770). Cell lines were validated by DNA sequencing (top panels). Data are representative of multiple experiments using fibroblasts from two independent FOP patients (GM00513 female 16 yrs and GM00783 male age unknown, Coriell Institute, NJ).

### Saracatinib induces dorsalization of zebrafish embryos

Dorsoventral axis specification during embryogenesis is regulated by the activity of BMP signaling agonists and antagonists (30). In fact, dorsalization of zebrafish embryos is a developmental phenotype that is highly specific for BMP inhibition and was the basis for the phenotypic screens that resulted in the identification of dorsomorphin (20). Incubating embryonic zebrafish with saracatinib starting at 0.5 hours post fertilization resulted in a dorsalized phenotype that increased in severity in a dose-dependent manner (Figure 4). We observed no evidence of cyclopia or phenotypes suggesting inhibition of mesoderm specification, consistent with the interpretation that saracatinib selectively inhibits BMP signaling in vivo without significantly impacting TGFβ or activin type I receptor activity.

**Figure 4.**
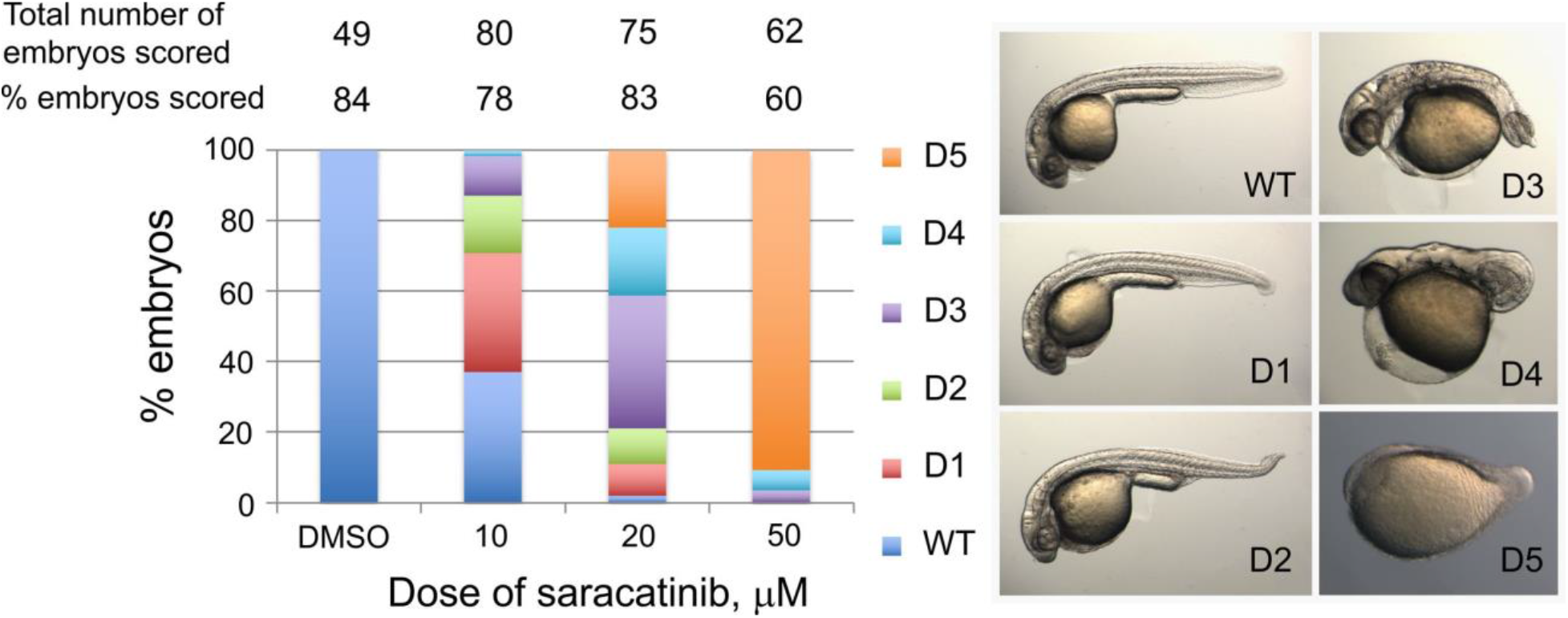
Saracatinib induces dorsalization of zebrafish embryos. Zebrafish embryos were treated from ≤ 0.5 hours post fertilization (hpf) with indicated concentrations of saracatinib delivered in 0.5% DMSO. Embryos were scored for dorsalization according to Mullins *et al*. (55). Representative embryos of each class are shown and reflect the increasing severity of dorsalization with increasing saracatinib concentration. WT-D4 images are of 30 hpf embryos, whereas the D5 embryo is shown at 12 hpf.

### Saracatinib prevents heterotopic bone formation in the *ACVR1^Q207D^*-Tg mouse model of HO

To evaluate the therapeutic potential of saracatinib for the treatment of FOP, we first employed one of the most studied models of HO, a Cre-inducible transgenic mouse that conditionally expresses the constitutively active *ACVR1^Q207D^* mutation (CAG-Z-eGFP-caALK2-Tg) (16, 31).

In these animals, activation of the *ACVR1^Q207D^* transgene and associated GFP reporter is mediated by a single intramuscular injection of Adenovirus expressing Cre recombinase (Ad.Cre) in the left hindlimb on postnatal day 7 (P7). This induces the expression of the human *ACVR1^Q207D^* transgene, as well as muscle necrosis and inflammation, which together result in the formation of heterotopic bone lesions within 7-10 days following Ad.Cre injection (16). This model is believed to recapitulate aspects of clinical HO and FOP in which muscle injury and inflammation potentiate the formation of heterotopic bone in soft tissues. Untreated, this process normally leads to progressive loss of passive and active range-of-motion of the hip, knee and ankle joints over 3-4 weeks, which is accompanied by the formation of intramuscular heterotopic bone lesions visible by X-ray (Figure 5).

**Figure 5.**
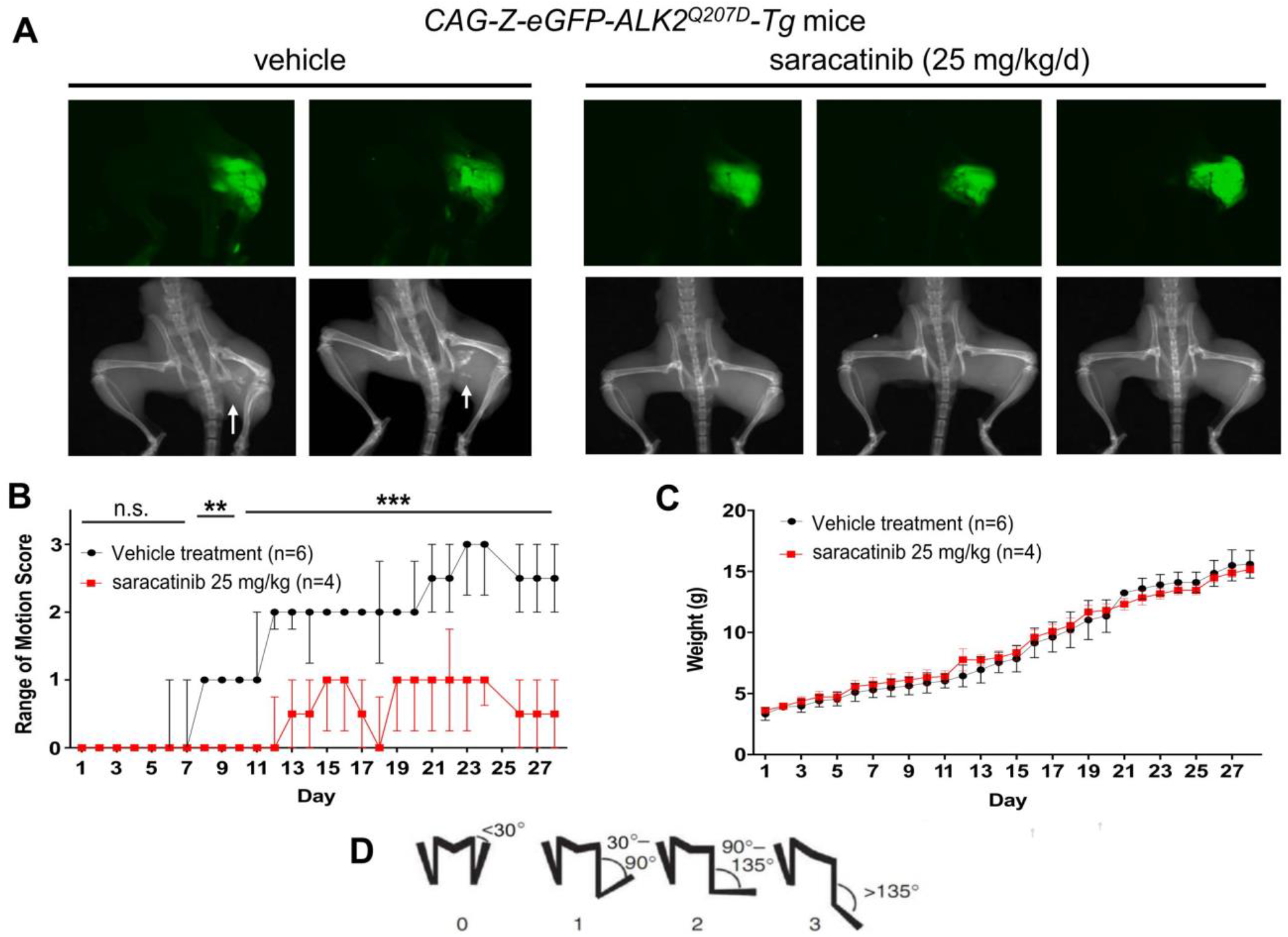
In vivo efficacy of saracatinib in the *ACVR1^Q207D^* transgenic mouse model of FOP. Neonatal *CAG-Z-eGFP-caALK2* transgenic mice were injected with Ad.Cre (1 x 10^8^ pfu i.m. P7) and treated with 25 mg/kg/d of saracatinib or vehicle orally for 28 days. (**A**) All mice expressed the eGFP reporter in the injected left hindlimb, and 100% of vehicle-treated mice (6/6) developed radiographic HO and severe loss of passive range of motion. (**B**) Treatment with saracatinib essentially abrogated radiographic HO in 4/4 mice, preserving range of motion (two-way ANOVA with Sidak’s test for multiple comparisons, p = n.s. for days 1-7, **p<0.01 for days 8-10, ***p<0.001 for days 11-28. All vs. vehicle treatment), data depicted as median ± I.Q.R, n as indicated. (**C**) Treatment with Saracatinib had no significant impact on normal growth based on weight gain as compared to controls (two-way ANOVA, with Dunnet’s test for multiple comparison, p=n.s.), data depicted as Mean ± S.E.M., n as indicated. (**D**) Representative hindlimb scoring system. All range of motion scoring was performed by two separate operators who were blinded to the animal’s treatment condition.

Protocols for saracatinib use in mice are well established from oncology studies and typically use a clinically relevant dose of 25 mg/kg/day (29, 32–35). CAG-Z-eGFP-caALK2-Tg mice treated similarly with saracatinib by oral gavage (25 mg/kg once daily for 28 days) demonstrated significantly improved range of motion, and markedly reduced HO at the site of Ad.Cre injection by X-ray compared to mice treated with vehicle control (Figure 5). GFP expression in the left hind limb confirmed efficient recombination of the *ACVR1^Q207D^* transgene at the injection site in all drug- and vehicle-treated animals (Figure 5A).

Notably, saracatinib did not impact the normal growth of the pups based on weight gain curves, suggesting that saracatinib can be well tolerated at doses effective for inhibiting HO (Figure 5C). By contrast, ALK2 inhibitor tool compounds LDN-193189 and LDN-212854 (6 mg/kg i.p. twice daily) (17), caused a 10-25% weight loss relative to vehicle-treated animals under these regimens (Supplemental Figure 7).

### Other SRC inhibitors are ineffective in blocking HO

Saracatinib has known activity against osteoclasts and bone turnover through SRC that could potentially contribute to the phenotypic response (35, 36). To investigate the potential for a SRC-mediated effect we tested vandetanib (AZD6474) as a control compound in the same *ACVR1^Q207D^*-Tg mouse model. Vandetanib is an analogous quinazoline-based Src kinase inhibitor that was approved for clinical use in advanced medullary thyroid cancer (28). While it has proven potency against SRC and other tyrosine kinases, including VEGFR2, EGFR and RET (37), vandetanib does not show any significant binding to ALK2 (Supplemental Figure 1C). When administered at 25 mg/kg i.p. twice daily for 28 days, vandetanib did not block HO formation in *ACVR1^Q207D^*-Tg mice following injection with Ad.Cre (Supplemental Figure 8), despite an exposure higher than previously shown to demonstrate in vivo efficacy against other target kinases in rodent models (38). Thus, saracatinib’s beneficial effects in the FOP mouse model were not from inhibition of SRC.

### Saracatinib prevents heterotopic bone formation in the *Acvr1^[R206H]FlEx/+^* mouse model of FOP

To test the impact of saracatinib in a more faithful model of FOP disease, we employed a more recently described conditional *Acvr1^R206H^* knock-in mouse, in which the *Acvr1^R206H^* mutant allele (*Acvr1^[R206H]FlEx/+1]^*) is conditionally expressed within the mouse *Acvr1* locus following Cre-loxP-mediated recombination (13). *Acvr1^[R206H]FlEx/+]^* knock-in mice were challenged with Ad.Cre (1 x 10^8^ pfu intramuscularly) on P7 to induce Cre-mediated recombination as well as muscle injury. For initial testing, mice were then treated daily with 25 mg/kg of saracatinib or vehicle by oral gavage for 28 days and monitored up to 90 days without further treatment. The rate of HO induction in this FOP mouse model based on X-ray or range-of-motion loss was notably reduced, consistent with the milder gain of function of *Acvr1^R206H^* compared with *ACVR1^Q207D^* (Figure 6A-B). Vehicle-treated animals showed the first evidence of decreased range of motion at day 13, which subsequently deteriorated up to 90 days, when abundant HO was detected (Figure 6A-B). In contrast, animals receiving an initial 28 day course of saracatinib were protected for the duration of the study and only developed mild loss of range of motion between 60 and 90 days after completing treatment (Figure 6A-B). These initial results confirmed that saracatinib attenuates HO mediated by dysregulated *Acvr1^R206H^* signaling in vivo and may be an effective transient therapy following acute trauma, as supported by the prolonged protection observed after treatment withdrawal and the lack of rebound ossification.

**Figure 6.**
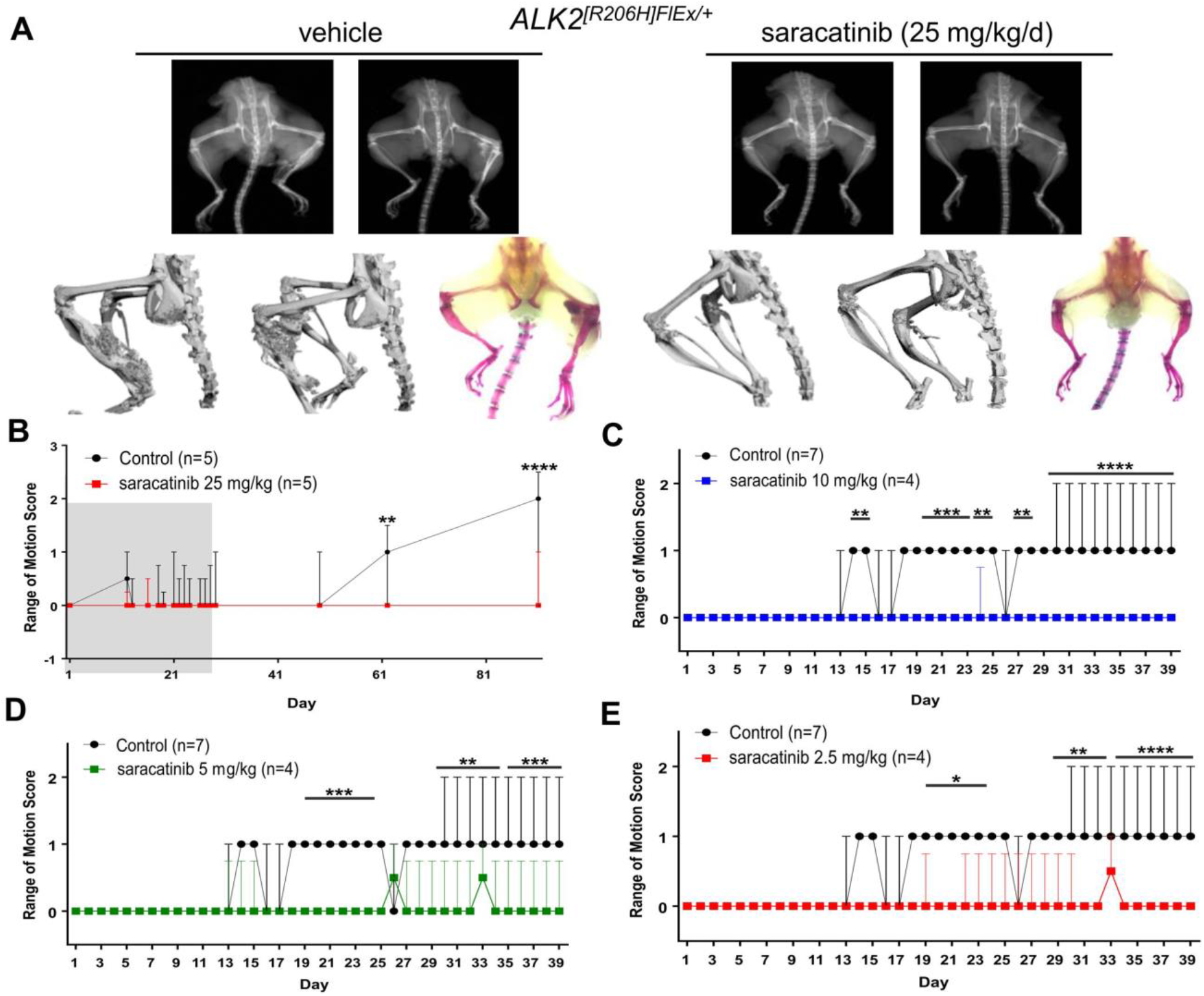
Dose dependent in vivo efficacy of saracatinib in the knock-in mouse model of FOP. Neonatal *Acvr1^[R206H]FlEX/+]^* mice were injected with a low dose of Ad.Cre (1 x 10^8^ pfu i.m. P7) and treated with 25 mg/kg/d of saracatinib or vehicle orally for 28 days (grey shaded region) and observed for a total of 90 days. Representative mice are shown with X-Ray Radiography, Micro-CT, and Alizarin red hindlimb prep. (**A**) Sixty percent of vehicle-treated mice (3/5) developed HO and severe loss of passive range of motion at 90 days. Treatment with saracatinib prevented radiographic HO in 5/5 treated mice, and (**B**) preserved range of motion in 4/5 mice at 90 days (two-way ANOVA with Sidak’s test for multiple comparisons, **p<0.01 at Day 61, ****p<0.0001 at Day 90 vs. control), data depicted as median ± I.Q.R., n as indicated. *ACVR1^[R206H]FlEX/+^* knock-in mice were injected with a high dose of Ad.Cre (5 x 10^9^ pfu i.m. P7) and treated with varying doses of saracatinib or vehicle orally for 40 days. Radiographic HO and associated impaired range of motion were observed to progress over 40 days of treatment. Treatment with saracatinib at 2.5, 5, and 10 mg/kg/d protected mice from radiographic HO, and preserved passive range of motion (**C-E**), (two-way ANOVA with Dunnet’s test for multiple comparisons, (C) *p<0.05 days 14-15,24-25,27-28, ***p<0.001 days 19-23, ****p<0.0001 days 29-39 vs. vehicle treatment; (**D**) ***p<0.001 days 19-24, **p<0.01 days 29-34, ****p<0.0001 days 35-39 vs. vehicle treatment; (**E**) *p<0.05 days 19-24, **p<0.01 days 29-33, ****p<0.0001 days 34-39 vs. vehicle treatment). Data depicted as median ± I.Q.R, n as indicated.

Chronic use of saracatinib as a SRC family inhibitor was recently explored in two phase II clinical studies in Alzheimer’s disease and lymphangioleiomyomatosis (ClinicalTrials.gov identifiers NCT02167256 and NCT02737202 respectively). Both studies used lower dosing regimens of 100-125 mg saracatinib daily for 9-12 months (4, 39, 40), corresponding to exposures of ~8 mg/kg/day orally in mice based on comparable levels of drug achieved in plasma and brain (41). Further experiments in the *ACVR1^[R206H]FlEx/+]^* knock-in mice were therefore performed using saracatinib in the range 2.5-10 mg/kg/day. A higher dose of Ad.Cre injection (5 x 10^9^ pfu) was administered to all animals to promote more rapid HO induction. Importantly, we observed that these reduced doses of saracatinib were also effective in preventing range-of-motion loss over a treatment course of 40 days (Figure 6C-E).

The efficacy observed at moderate doses of AZD0530 did not appear to impact neonatal growth assessed by weight gain, femur length, and bone mineral density (Figure 7). Thus, saracatinib may be sufficiently potent and well-tolerated for long-term administration in FOP, and in this manner could suppress acute flares as well as disease that progresses without known flares or antecedent injury.

**Figure 7.**
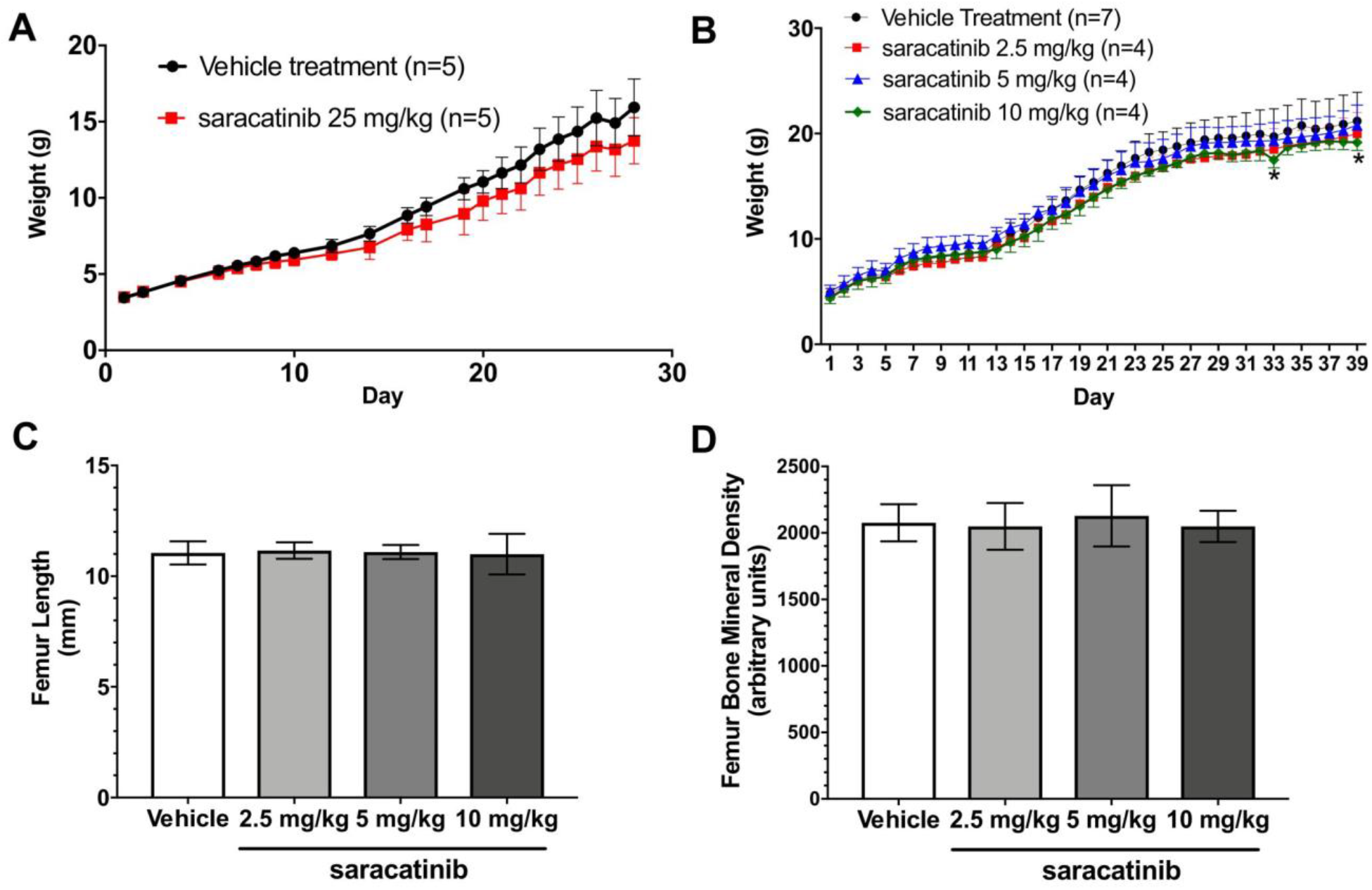
Saracatinib treatment preserves neonatal and juvenile orthotopic skeletal growth. (**A**) Treatment of neonatal mice with saracatinib at 25 mg/kg/d exerted a mild impact on weight gain during the first 28 days of treatment (p<0.0001 compared to vehicle treatment, two-way ANOVA), data depicted as mean ± S.E.M., n as indicated. (**B**) Treatment of neonatal mice with saracatinib at 2.5 – 10 mg/kg/d did not impact weight gain (p=NS compared to vehicle treatment, two-way ANOVA), nor did it impact neonatal and juvenile orthotopic skeletal growth, based on preserved femur length (**C**), and preserved femur bone mineral density (**D**) at 2.5, 5, and 10 mg/kg/d oral doses of saracatinib versus vehicle treatment (data shown as mean ± S.D., n=8, 4, and 4 femurs analysed respectively; one-way ANOVA with Dunnett’s test for multiple comparisons).

## Discussion

Saracatinib (AZD0530) is a potent, orally bioavailable SRC/ABL inhibitor originally developed by AstraZeneca for treatment of ovarian adenocarcinoma (29). Saracatinib exhibits excellent pharmacokinetic parameters (t_1/2_~40 hours) and is safe in healthy volunteers up to a daily dose of 250 mg (36). Despite extensive testing in the clinic, with >30 registered trials in more than 600 patients, saracatinib has shown only limited efficacy in cancer therapy. However, saracatinib has proven to be of significant interest for drug repositioning studies and now forms part of an open-innovation portfolio of industry assets made available to the US National Institutes of Health and the Medical Research Council in the UK (4).

Based on a screen of known biologically active compounds that are approved or in clinical testing, we identified saracatinib as a multi-kinase inhibitor that potently inhibits ALK2 in addition to the SRC family of human kinases. Within the BMP/TGFβ type I receptor family, saracatinib exhibits useful selectivity; it discriminates between the signalling of ALK2 and its cognate ligands in wild-type cells (BMP6 and BMP7) and mutant cells (activin A) versus other receptors and ligands of this family. In fact, saracatinib and the previously characterized pyrazolo-pyrimidine compound LDN-193189 display comparable cellular potency against caALK2 (IC_50_ values of 14 nM and 11 nM respectively), as well as selectivity against caALK5 (IC_50_ values of 217 nM and 213 nM respectively) (17). The ability of saracatinib to inhibit ALK2 signalling in a selective manner in vivo was confirmed by its ability to induce dorsalization in embryonic zebrafish, without inducing other phenotypes due to the suppression of TGFβ or activin signalling.

Historically, much drug discovery effort has focused on tyrosine kinases of relevance to cancer (28). The BMP/TGFβ receptors form the only family of serine/threonine transmembrane receptor kinases but map favourably for drug repositioning to the tyrosine kinase-like (TKL) branch of the human kinome (42). ALK2 therefore shares significant sequence and structural similarity with the tyrosine kinases, including a conserved gatekeeper threonine residue in the hinge region of the ATP-binding pocket (14). In contrast, our screens identify little saracatinib activity against most other serine/threonine kinases. This specificity appears to derive from the large chlorobenzodioxole moiety of saracatinib which complements the size and shape of the back pocket in ALK2. Notably, these regions differ in the co-structures of ALK2 and SRC due to the different positions of the αC helix in their inactive states. Based on these differences, it is possible that further optimization of this chemotype could yield molecules that bind with increased selectivity for ALK2 over SRC family kinases.

Importantly, our findings demonstrate that when FOP is modelled in an authentic and physiologic manner by expressing the classic FOP-causing *ACVR1^R206H^* allele under endogenous promoter control, short-term dosing of saracatinib is sufficient to suppress the development of HO for an extended period following injury. One might extrapolate from these findings that acute prophylactic pharmacologic inhibition of ALK2 activity could be sufficient to prevent HO in individuals with FOP, although the specific window of time during which transient therapy is effective would need to be determined in man. BMP receptor signalling has also been implicated in the formation of HO in the general population without *ACVR1* mutation, including burn- and trauma-associated HO, or HO associated with severe neurologic injuries (43, 44). Thus, saracatinib is an interesting clinical candidate to test as an acute therapy in these disease models as well as in FOP patients.

While several classes of ALK2 kinase inhibitors have been reported in the literature previously, saracatinib has potential advantages for clinical translation due to the detailed characterization of its tolerability and side-effect profile in man. In the case of FOP, the high risk and large investment associated with developing a novel compound, coupled with the extremely small numbers of patients eligible for trials and treatment could further hinder drug development efforts. In addition to being used in numerous oncology clinical trials, saracatinib has been explored as a FYN kinase inhibitor in a phase II study of patients with mild-to-moderate Alzheimer’s disease at doses up to 125 mg daily for 12 months (4, 39, 40) (NCT02167256). Another phase II study is investigating this molecule as a SRC inhibitor in lymphangioleiomyomatosis with a regimen of 125 mg daily for 9 months (4) (NCT02737202). The primary side effect of saracatinib at these doses has been gastrointestinal discomfort (40). The finding that saracatinib is tolerable when administered chronically at moderate doses suggests that its application to FOP could be extended beyond the prophylactic treatment of acute episodes of HO. It has been generally supposed that much of FOP disease progression occurs episodically, in acute flares characterized by soft tissue swelling, pain, and warmth (45). Indeed, phase II clinical trials for FOP using the RARγ agonist palovarotene were designed to treat acute flare episodes in an abortive manner (NCT02190747). Recent natural history studies have suggested, however, that nearly half of individuals with FOP report the insidious progression of disease without known flares or antecedent injury (46). Thus, more chronic suppression of the underlying disease process outside of acute flares may be needed. We observed that reduced doses of saracatinib (2.5-10 mg/kg/day) that were similar to or lower than exposures in phase II studies in Alzheimer’s disease and lymphangioleiomyomatosis were effective in preventing HO in the *ACVR1^R206H^* FOP mouse model. Thus, if saracatinib is sufficiently tolerable, its chronic use for the long-term suppression of *ACVR1^R206H^* activity might be a promising option for maintaining optimal control of FOP disease, suppressing its progression even in the absence of flares or acute episodes, and potentially preventing flares altogether. Based on the evidence of target engagement and efficacy in pre-clinical models at doses equivalent to or lower than those previously tolerated in humans, a phase II clinical trial using 100 mg/day has been launched to explore the use of saracatinib as prophylaxis for FOP progression (NCT04307953).

The discovery that all cases of FOP are caused by gain of function mutations in the intracellular domain of ALK2/*ACVR1* has ignited great interest in ALK2 as a therapeutic target. Repositioning saracatinib for FOP offers an accelerated path to clinical proof of efficacy studies and potentially significant benefits to individuals with this devastating condition.

## Methods

### Protein expression and purification

All constructs for protein expression were cloned into the pFastBac-derived vector pFB-LIC-Bse, which provides an N-terminal hexahistidine tag to enable purification. For DSF, constructs included ALK2 GS and kinase domain residues 172-499 (wild type or R206H) or 172-509 (L196P, Q207E, R258S, or G328E), while the ALK5 construct included residues 162-503. For crystallography, the ALK2 construct comprised residues 201-499 with the Q207D mutation. Baculoviral expression was performed in Sf9 insect cells at 27°C, shaking at 110 rpm. Cells were harvested at 72 hours post infection and resuspended in 50 mM HEPES pH 7.5, 500 mM NaCl, 5 mM imidazole, 5% glycerol, 0.1 mM Tris(2-carboxyethyl)phosphine (TCEP), supplemented with protease inhibitor set V (Calbiochem). Cells were lysed using sonication (Vibra-cell). Subsequently, 0.15% polyethylenimine was added to precipitate DNA before the insoluble material was excluded by centrifugation at 21,000 rpm. Initial purifications were performed by Ni-affinity chromatography. Eluted proteins were cleaved with Tobacco Etch Virus (TEV) protease and further purified by size exclusion chromatography using a S200 HiLoad 16/60 Superdex column buffered in 50 mM HEPES pH 7.5, 300 mM NaCl, 1 mM TCEP. Excess protein was flash frozen and stored at −80°C.

### Structure Determination

Frozen ALK2 protein was thawed, purified from aggregates on a S200 HiLoad 16/60 Superdex column and concentrated to 10 mg/mL buffered in 5 mM HEPES pH 7.5, 100 mM NaCl. Crystallization was performed using the sitting drop vapour diffusion method at 4°C. Viable crystals of ALK2 in complex with saracatinib grew in a 150 nL drop mixing 10 mg/mL protein, pre-incubated with 1 mM compound, with a reservoir solution containing a 1:1 volume ratio mix of 1.26 M sodium phosphate monobasic and 0.14 M potassium phosphate dibasic. Crystals were transferred into a cryo-protective solution prepared from the mother liquor supplemented with 25% ethylene glycol. Diffraction data were collected at Diamond Light Source, beamline I04-1 at a temperature of 100 K using a wavelength of 0.9174 Å. Data were processed and scaled with MOSFLM and SCALA from CCP4 suite (47, 48). The structure was solved by molecular replacement using PHASER (49) and the co-ordinates of ALK2 from the ALK2-FKBP12-dorsomorphin complex (14) as a search model. Subsequent manual model building was performed using COOT (50) alternated with refinement in REFMAC (51) and Phenix (52). TLS-restrained refinement was applied in the latter cycles using the input thermal motion parameters determined by the TLSMD server (53). The final model was verified for geometry correctness with MOLPROBITY (54) and deposited in the Protein Databank (PDB accession 6ZGC). The Ramachandran statistics after refinement were 99.8% preferred and 98.0% allowed. Data collection and refinement statistics are summarized in Supplemental Table 2.

### DSF screening

Differential scanning fluorimetry (DSF) was performed as described by Fedorov et al. (10). Purified proteins containing the GS and kinase domains of ALK2 or ALK5 were screened at 2 μM concentration against a library of ~150 clinically tested kinase inhibitors (Selleckchem) in a 96-well plate format. Compounds were added to a final concentration of 12.5 μM in an assay buffer containing 10 mM HEPES pH 7.4, 150 mM NaCl and SYPRO Orange (1:1000 dilution, Sigma). Samples were heated from 25 to 96°C in a Mx3005P real-time PCR instrument (Stratagene). Fluorescence was monitored with excitation and emission filters set to 465 and 590 nm, respectively. Data were analyzed with the MxPro software. Thermal shift values (ΔTm) induced by inhibitor binding were calculated relative to control wells containing protein and 2.5% DMSO.

### Recombinant ALK1-6 enzyme assays

Recombinant ALK1 was obtained from Life Technology (catalog# PV4883). Recombinant ALK2 to ALK6 were purchased from Carna Bioscience (catalog# 09-134, Natick, MA, USA). A LANCE Ultra TR-FRET kinase assay was used for measurements of enzyme activities that included an-Ulight labeled (Thr-1342) peptide substrate (catalog #TRF0130-M, PerkinElmer) and a Europium-labeled anti-phospho peptide antibody (catalog # TRF0218-M). The kinase reaction was carried out in a buffer containing 10 mM MgCl_2_, 3 mM MnCl_2_, 0.005% Tween-20, 2 mM DTT, and 50 mM HEPES pH 7.0. The details of the enzyme assays will be reported in a separate paper (Huang X. et al. manuscript in preparation). Briefly, 2.5 μL/well of kinase protein (10 nM final) was dispensed to a white solid 1536-well plate followed by addition of 23 nL/well of compound diluted in DMSO using a Pin-tool station (Wako-Kalypsis). After a 10-min incubation at room temperature, 2.5 μL/well of substrate solution (50 μM peptide and 10 μM ATP, final) was added and the assay plate was incubated for 60 min at room temperature. The enzyme reaction was stopped by addition of 5 μL/well of detection solution containing the Europium-labeled anti-phospho peptide antibody. The signals of time-resolved fluorescence resonance energy transfer (TR-FRET) assay were measured in an Envision plate reader (PerkinElmer) with a TR-FRET mode (Ex=320, Em1=665, Em2=615 nm).

### Cell viability

Cell viability was assayed with an MTT (3-(4,5-dimethylthiazol-2-yl)-2,5-diphenyl tetrazolium bromide) colorimetric assay (Promega) per manufacturer’s instructions.

### Western blotting

Bovine aortic endothelial cells (BAEC) were purchased from Lonza and maintained in high glucose DMEM supplemented with 10% fetal bovine serum (FBS). C2C12 myofibroblast cells from the European Collection of Cell Cultures (ECACC) were grown in DMEM/F12 with 10% FCS. Primary human fibroblasts GM00513 (FOP R206H), GM00783 (FOP R206H) and ND34770 (wild type) were obtained from Coriell (ccr.coriell.org/ninds) and grown in MEM with 2 mM L-glutamine and 10% FCS. Western blotting was performed essentially as described in Sanvitale *et al* (26). Briefly, 1 x 10^5^ cells were seeded into 6-well plates. The following day cells were starved in medium containing 1% FCS for 5 hours before treatment with indicated concentrations of saracatinib and growth factor for 1 hour. Cells were lysed, proteins separated by SDS-PAGE and analyzed by Western blotting using relevant antibodies (Cell Signaling: anti-SMAD1 #9743), anti-P-SMAD1/5 #9516, anti-SMAD2 #5339, anti-P-SMAD2 #3101).

### In-Cell Immunofluorescent Assay of phospho-SMAD1/5 levels

Human FOP fibroblasts (Coriell Institute, NJ) were grown to confluence in DMEM supplemented with 10% FBS (Sigma-Aldrich, MO) and then seeded into 96-well plates (Costar^®^ 3610; Corning, NY). The cells were incubated at 37°C & 10% CO_2_, and allowed to attach; following this, they were starved overnight in 1% FBS-DMEM. Saracatinib (Biotang Inc., MA) was dissolved in DMSO (Fisher Scientific, NH), while LDN-193189 (Yu Lab, MA) was diluted in 0.1% PBS-BSA. The cells were then pre-incubated with increasing molar concentrations of saracatinib and LDN-193189 for 15 minutes. Activin A (R&D Systems, MN) was diluted in 0.1% PBS-BSA (Sigma-Aldrich, MO) to a working concentration of 250 ng/mL. After addition of activin A, the cells were allowed to incubate for 45 minutes. Fixation of the cells was performed with ice cold methanol and 0.5% glutaraldehyde (Fisher Scientific, NH), before the addition of primary antibody (Phospho-Smad1/5 (Ser463/465) Rabbit mAb #9516; Cell Signaling Technologies, MA) and secondary antibody (Anti-rabbit IgG, HRP-linked antibody; Cell Signaling Technologies, MA). Finally, the cells were developed with BioFX Chemiluminescent Ultra Sensitive HRP Microwell Substrate solution (Surmodics, MN) and read on a Spectra Max L plate reader (Molecular Devices, CA). The software Softmax Pro was used, on the Ready to Glow Luciferase Assay setting with an integration time at 0.25 seconds and a wavelength of 470 nm.

### Dual luciferase assays of constitutively active receptors

C2C12 cells stably expressing firefly luciferase under the control of BMP-responsive promoter element (BRE-Luc) were generously provided by Dr. Peter ten Dijke (Leiden University Medical Center, NL). Human embryonic kidney (HEK) 293T cells stably transfected with the TGFβ-responsive element fused to luciferase gene (CAGA-Luc) were a kind gift of Dr. Howard Weiner (Brigham and Women’s Hospital, Boston, MA). C2C12 BRE-Luc and 293T CAGA-Luc cells were seeded at 20,000 cells in DMEM supplemented with 2% FBS per well in tissue culture treated 96-well plates (Costar^®^ 3610; Corning). The cells were incubated for 1 hour (37°C and 10% CO_2_) and allowed to settle and attach. Saracatinib was diluted in DMSO, and diluted drug or DMSO vehicle only added to cells at final compound concentrations of 1 nM to 10 μM and a final concentration of DMSO of 2%. Cells were then incubated for 30 min. Adenovirus expressing constitutively active BMP and TGFβ type I receptors (Ad.caALK1-5), generously provided by Dr. Akiko Hata (University of California at San Francisco), were added to achieve a multiplicity of infection (MOI) of 100. Plates were incubated overnight at 37°C. Media was discarded and firefly luciferase activity was measured (Promega) according to manufacturer’s protocol. Light output was measured using a Spectramax L luminometer (Molecular Devices) with an integration time of one second per well. Data were normalized to 100% of incremental BRE-Luc activity due to adenoviruses specifying caALK1, 2, or 3, or the incremental CAGA-Luc activity due to adenoviruses specifying caALK4 or 5, and also normalized to total cell counts based on MTT cell viability assay. Graphing and regression analysis by sigmoidal doseresponse with variable Hill coefficient were performed using GraphPad Prism software.

### Dual luciferase assays of ligand-induced signalling

MDA-MB-231 cells stably expressing BRE-Luc or CAGA-Luc were generously provided by Dr. Caroline Hill (Francis Crick Institute, UK). Some 4000 cells were seeded into each well of a 96-well plate in DMEM/F12 with 10% FCS. The following day the medium was replaced with DMEM/F12 and 1% FCS and then saracatinib added at the indicated concentrations to 6 wells, followed by overnight stimulation with growth factor. Luciferase activity was measured using the Promega Dual Luciferase Reporter Assay system and percentage activity determined compared to cells treated with growth factor and no inhibitor. Growth factors (Peprotech) were used at the following concentrations: BMP2 500 ng/mL, BMP4 10 ng/mL, BMP6 500 ng/mL, BMP7 500 ng/mL, TGFβ 4 ng/mL, activin A 200 ng/mL. Experiments were repeated at least 3 times and plotted as mean +/- SD.

### Zebrafish Dorsalization

Zebrafish embryos were treated with varying inhibitor concentrations from ≤ 0.5 hours post fertilization (hpf). Inhibitor stocks (20 mM) were diluted in fish water from DMSO. Embryos were scored according to Mullins *et al* (55).

### *ACVR1^Q207D^*-Tg and *Acvr1^[R206H]FlEx/+^* mouse models of heterotopic ossification

Mice were maintained in accordance with Institutional Animal Care and Use Committee guidelines under approved experimental protocols. Cre-inducible *ACVR1^Q207D^* (CAG-Z-eGFP-caACVR1-Tg) transgenic mice were a generous gift from Dr. Yuji Michina (University of Michigan, Ann Harbor, MI) as previously described (31). *Acvr1^[R206H]FlEX/+^* mice were generated as previously described (13). Heterotopic bone formation in these mice was introduced via single retro-popliteal injection of adenoviral Cre-recombinase at 1×10^8^ PFU at postnatal day 7 (P7). For initial experiments mice (n=3-4 per group) were treated once a day for 4 weeks with saracatinib at 25 mg/kg by oral gavage dissolved in a vehicle consisting of 5% DMSO and 95% peanut oil, or vehicle alone. Bone formation as a function of a loss of passive range of motion, via dorsiflexion of the left ankle joint was assessed daily. Scores were assessed by two independent observers blinded to genotype and treatment. The observers scored the minimum angle formed by the ankle and the tibia with passive dorsiflexion under light manual pressure as follows: 0, normal flexion with a minimal angle of < 30°; 1, mildly impaired flexion with a minimal angle of ≥ 30° but < 90°; 2, moderately impaired flexion with a minimal angle of ≥ 90° but < 135°; and 3, severely impaired flexion with a minimal angle of ≥ 135° (depicted in Figure 5D).

To monitor the progression of heterotopic ossification during the in-vivo studies, X-ray radiographs were also imaged at various time points. Mice were anesthetized using ketamine at 100 mg/kg, and imaged non-terminally in the prone position with tibial lateralization and hip abduction, allowing for reproducible positioning. Radiographs were obtained using 30 second scans and 50 kV exposures (MS FX In-Vivo Pro: CareSteam Imaging, Rochester NY). Heterotopic ossification was additionally examined with micro-computed tomography (micro-CT instrument μCT35, ScanCo, 70 kV, 50 mA, 32 millisecond exposure, 220 views per rotation, 0.877 increment angle). Mice were anesthetized with ketamine at 125 mg/kg, limbs slightly restrained with tape, and imaged non-terminally in the supine position. After reconstruction, regions of interest were chosen and exported into DICOM format, and analysed by Osirix analysis software.. Fixed soft tissues and bone were stained with Alizarin red and Alcian blue as previously described (56). Further studies of saracatinib dose response relationships were performed in the *Acvr1^R206H^* mice challenged with Ad.Cre (5 x 10^9^ pfu) as above and then treated daily with either 10, 5, 2.5 or 0 mg/kg of saracatinib by oral gavage for 40 days. All experiments were performed using mixed gender populations with no observed gender bias in outcomes.

### Statistics

Data are expressed as mean ± SEM unless stated otherwise. A Student’s t test 2-tailed was used where indicated. A P value <0.05 was considered significant.

### Study approval

Mice were maintained in accordance with Harvard Medical School and Brigham and Women’s Hospital Institutional Animal Care and Use Committee guidelines under approved experimental protocols.

## Competing financial interests

A.N.E. is an employee of Regeneron Pharmaceuticals, Inc. P.B.Y. is a co-founder and holds stock in Keros Therapeutics, which develops therapies for hematologic and musculoskeletal diseases targeting BMP and TGFβ signaling pathways, including FOP. P.B.Y.’s interests are reviewed and managed by Brigham and Women’s Hospital in accordance with their conflict of interest policies. The remaining authors have no financial disclosures.

For publication under a Creative Commons CC-BY license

## Author contributions

A.N.B. and P.B.Y. designed the research. E.W., J.B., G.K., E.S.P. and P.B.Y. performed research and analyzed the data. D.D., A.H.M., Y.S., G.A.B., X.H., P.E.S. and W.Z. contributed additional experimental data. E.W, J.B., G.K., E.S.P., A.N.E., J.C.S., P.B.Y. and A.N.B wrote and revised the paper.

## Acknowledgements

The authors would like to thank Diamond Light Source for beamtime (proposal mx10619), as well as the staff of beamlines I03 and I04-1 for assistance with crystal testing and data collection. This work was supported by US National Institutes of Health Grants AR057374 (P.B.Y.) and HL007604 (BWH Cardiovascular Division T32 supporting D.D. and J.B.), Department of Defense Grant MR140072 (P.B.Y.), a Harvard Stem Cell Institute Seed Award (P.B.Y.), a Leducq Foundation Transatlantic Network of Excellence Award (P.B.Y. and J.C.S.), the Francis Crick Institute, which receives its core funding from Cancer Research UK (FC001-157), the UK Medical Research Council (FC001-157), and Wellcome (FC001-157) (J.C.S.), a Howard Hughes Medical Institute Early Career Physician-Scientist Award (P.B.Y.), and from the International FOP Association (P.B.Y., D.D.X. and Y.S.). E.W., G.K. and A.N.B. acknowledge support from the SGC, a registered charity (number 1097737) that receives funds from AbbVie, Bayer Pharma AG, Boehringer Ingelheim, Canada Foundation for Innovation, Eshelman Institute for Innovation, Genentech, Genome Canada through Ontario Genomics Institute [OGI-196], EU/EFPIA/OICR/McGill/KTH/Diamond Innovative Medicines Initiative 2 Joint Undertaking (EUbOPEN grant number 875510), Janssen, Merck & Co., MSD, Novartis Pharma AG, Pfizer, São Paulo Research Foundation-FAPESP, Takeda and Wellcome [106169/ZZ14/Z]. A.N.B. and P.B.Y. acknowledge support from Innovative Medicines Initiative 2 Joint Undertaking (STOPFOP grant number 821600).

## References

1. Boycott KM, Vanstone MR, Bulman DE, and MacKenzie AE. Rare-disease genetics in the era of next-generation sequencing: discovery to translation. Nature reviews Genetics. 2013;14(10):681–91.

2. Melnikova I. Rare diseases and orphan drugs. Nat Rev Drug Discov. 2012;11(4):267–8.

3. Braun MM, Farag-El-Massah S, Xu K, and Cote TR. Emergence of orphan drugs in the United States: a quantitative assessment of the first 25 years. Nat Rev Drug Discov. 2010;9(7):519–22.

4. Frail DE, Brady M, Escott KJ, Holt A, Sanganee HJ, Pangalos MN, et al. Pioneering government-sponsored drug repositioning collaborations: progress and learning. Nat Rev Drug Discov. 2015;14(12):833–41.

5. Ekins S, Williams AJ, Krasowski MD, and Freundlich JS. In silico repositioning of approved drugs for rare and neglected diseases. Drug Discov Today. 2011;16(7–8):298–310.

6. Nosengo N. Can you teach old drugs new tricks? Nature. 2016;534(7607):314–6.

7. Boycott KM, Dyment DA, Sawyer SL, Vanstone MR, and Beaulieu CL. Identification of genes for childhood heritable diseases. Annu Rev Med. 2014;65:19–31.

8. Kaplan FS, Glaser DL, Pignolo RJ, and Shore EM. A new era for fibrodysplasia ossificans progressiva: a druggable target for the second skeleton. Expert Opinion on Biological Therapy. 2007;7(5):705–12.

9. Kaplan FS, Zasloff MA, Kitterman JA, Shore EM, Hong CC, and Rocke DM. Early mortality and cardiorespiratory failure in patients with fibrodysplasia ossificans progressiva. J Bone Joint Surg Am. 2010;92(3):686–91.

10. Dey D, Bagarova J, Hatsell SJ, Armstrong KA, Huang L, Ermann J, et al. Two tissue-resident progenitor lineages drive distinct phenotypes of heterotopic ossification. Sci Transl Med. 2016;8(366):366ra163.

11. Shore EM, Xu M, Feldman GJ, Fenstermacher DA, Cho TJ, Choi IH, et al. A recurrent mutation in the BMP type I receptor ACVR1 causes inherited and sporadic fibrodysplasia ossificans progressiva. Nat Genet. 2006;38(5):525–7.

12. Kaplan FS, Xu M, Seemann P, Connor JM, Glaser DL, Carroll L, et al. Classic and atypical fibrodysplasia ossificans progressiva (FOP) phenotypes are caused by mutations in the bone morphogenetic protein (BMP) type I receptor ACVR1. Human Mutation. 2009;30(3):379–90.

13. Hatsell SJ, Idone V, Wolken DM, Huang L, Kim HJ, Wang L, et al. ACVR1R206H receptor mutation causes fibrodysplasia ossificans progressiva by imparting responsiveness to activin A. Sci Transl Med. 2015;7(303):303ra137.

14. Chaikuad A, Alfano I, Kerr G, Sanvitale CE, Boergermann JH, Triffitt JT, et al. Structure of the Bone Morphogenetic Protein Receptor ALK2 and Implications for Fibrodysplasia Ossificans Progressiva. Journal of Biological Chemistry. 2012;287(44):36990–8.

15. Hino K, Ikeya M, Horigome K, Matsumoto Y, Ebise H, Nishio M, et al. Neofunction of ACVR1 in fibrodysplasia ossificans progressiva. Proc Natl Acad Sci U S A. 2015;112(50):15438–43.

16. Yu PB, Deng DY, Lai CS, Hong CC, Cuny GD, Bouxsein ML, et al. BMP type I receptor inhibition reduces heterotopic ossification. Nature Medicine. 2008;14(12):1363–9.

17. Mohedas AH, Xing X, Armstrong KA, Bullock AN, Cuny GD, and Yu PB. Development of an ALK2-Biased BMP Type I Receptor Kinase Inhibitor. ACS Chemical Biology. 2013;8(6):1291–302.

18. Agarwal S, Loder S, Brownley C, Cholok D, Mangiavini L, Li J, et al. Inhibition of Hif1alpha prevents both trauma-induced and genetic heterotopic ossification. Proc Natl Acad Sci U S A. 2016;113(3):E338–47.

19. Shimono K, Tung WE, Macolino C, Chi AH, Didizian JH, Mundy C, et al. Potent inhibition of heterotopic ossification by nuclear retinoic acid receptor-gamma agonists. Nat Med. 2011;17(4):454–60.

20. Yu PB, Hong CC, Sachidanandan C, Babitt JL, Deng DY, Hoyng SA, et al. Dorsomorphin inhibits BMP signals required for embryogenesis and iron metabolism. Nat Chem Biol. 2008;4(1):33–41.

21. Mohedas AH, Wang Y, Sanvitale CE, Canning P, Choi S, Xing X, et al. Structure-activity relationship of 3,5-diaryl-2-aminopyridine ALK2 inhibitors reveals unaltered binding affinity for fibrodysplasia ossificans progressiva causing mutants. J Med Chem. 2014;57(19):7900–15.

22. Engers DW, Frist AY, Lindsley CW, Hong CC, and Hopkins CR. Synthesis and structure-activity relationships of a novel and selective bone morphogenetic protein receptor (BMP) inhibitor derived from the pyrazolo[1.5-a]pyrimidine scaffold of dorsomorphin: the discovery of ML347 as an ALK2 versus ALK3 selective MLPCN probe. Bioorg Med Chem Lett. 2013;23(11):3248–52.

23. Fabbro D, Cowan-Jacob SW, Mobitz H, and Martiny-Baron G. Targeting cancer with small-molecular-weight kinase inhibitors. Methods Mol Biol. 2012;795:1–34.

24. Wu P, Nielsen TE, and Clausen MH. Small-molecule kinase inhibitors: an analysis of FDA-approved drugs. Drug Discov Today. 2016;21(1):5–10.

25. Fedorov O, Niesen FH, and Knapp S. Kinase inhibitor selectivity profiling using differential scanning fluorimetry. Methods Mol Biol. 2012;795:109–18.

26. Sanvitale CE, Kerr G, Chaikuad A, Ramel MC, Mohedas AH, Reichert S, et al. A new class of small molecule inhibitor of BMP signaling. PLoS One. 2013;8(4):e62721.

27. Cuny GD, Yu PB, Laha JK, Xing X, Liu J-F, Lai CS, et al. Structure-activity relationship study of bone morphogenetic protein (BMP) signaling inhibitors. Bioorganic & Medicinal Chemistry Letters. 2008;18(15):4388–92.

28. Wu P, Nielsen TE, and Clausen MH. FDA-approved small-molecule kinase inhibitors. Trends Pharmacol Sci. 2015;36(7):422–39.

29. Hennequin LF, Allen J, Breed J, Curwen J, Fennell M, Green TP, et al. N-(5-chloro-1,3-benzodioxol-4-yl)-7-[2-(4-methylpiperazin-1-yl)ethoxy]-5-(tetrahydro-2H-pyran-4-yloxy)quinazolin-4-amine, a novel, highly selective, orally available, dual-specific c-Src/Abl kinase inhibitor. J Med Chem. 2006;49(22):6465–88.

30. Wu MY, and Hill CS. TGF-beta Superfamily Signaling in Embryonic Development and Homeostasis. Developmental Cell. 2009;16(3):329–43.

31. Fukuda T, Scott G, Komatsu Y, Araya R, Kawano M, Ray MK, et al. Generation of a mouse with conditionally activated signaling through the BMP receptor, ALK2. Genesis. 2006;44(4):159–67.

32. Chang YM, Bai L, Liu S, Yang JC, Kung HJ, and Evans CP. Src family kinase oncogenic potential and pathways in prostate cancer as revealed by AZD0530. Oncogene. 2008;27(49):6365–75.

33. Green TP, Fennell M, Whittaker R, Curwen J, Jacobs V, Allen J, et al. Preclinical anticancer activity of the potent, oral Src inhibitor AZD0530. Mol Oncol. 2009;3(3):248–61.

34. Cavalloni G, Peraldo-Neia C, Sarotto I, Gammaitoni L, Migliardi G, Soster M, et al. Antitumor activity of Src inhibitor saracatinib (AZD-0530) in preclinical models of biliary tract carcinomas. Mol Cancer Ther. 2012;11(7):1528–38.

35. Heusschen R, Muller J, Binsfeld M, Marty C, Plougonven E, Dubois S, et al. SRC kinase inhibition with saracatinib limits the development of osteolytic bone disease in multiple myeloma. Oncotarget. 2016.

36. Hannon RA, Clack G, Rimmer M, Swaisland A, Lockton JA, Finkelman RD, et al. Effects of the Src kinase inhibitor saracatinib (AZD0530) on bone turnover in healthy men: a randomized, double-blind, placebo-controlled, multiple-ascending-dose phase I trial. J Bone Miner Res. 2010;25(3):463–71.

37. Jia HY, Wu JX, Zhu XF, Chen JM, Yang SP, Yan HJ, et al. ZD6474 inhibits Src kinase leading to apoptosis of imatinib-resistant K562 cells. Leuk Res. 2009;33(11):1512–9.

38. Gule MK, Chen Y, Sano D, Frederick MJ, Zhou G, Zhao M, et al. Targeted therapy of VEGFR2 and EGFR significantly inhibits growth of anaplastic thyroid cancer in an orthotopic murine model. Clin Cancer Res. 2011;17(8):2281–91.

39. Nygaard HB, Wagner AF, Bowen GS, Good SP, MacAvoy MG, Strittmatter KA, et al. A phase Ib multiple ascending dose study of the safety, tolerability, and central nervous system availability of AZD0530 (saracatinib) in Alzheimer’s disease. Alzheimers Res Ther. 2015;7(1):35.

40. van Dyck CH, Nygaard HB, Chen K, Donohue MC, Raman R, Rissman RA, et al. Effect of AZD0530 on Cerebral Metabolic Decline in Alzheimer Disease: A Randomized Clinical Trial. JAMA Neurol. 2019.

41. Kaufman AC, Salazar SV, Haas LT, Yang J, Kostylev MA, Jeng AT, et al. Fyn inhibition rescues established memory and synapse loss in Alzheimer mice. Ann Neurol. 2015;77(6):953–71.

42. Manning G, Whyte DB, Martinez R, Hunter T, and Sudarsanam S. The protein kinase complement of the human genome. Science. 2002;298(5600):1912–34.

43. Peterson JR, De La Rosa S, Eboda O, Cilwa KE, Agarwal S, Buchman SR, et al. Treatment of heterotopic ossification through remote ATP hydrolysis. Sci Transl Med. 2014;6(255):255ra132.

44. Qureshi AT, Crump EK, Pavey GJ, Hope DN, Forsberg JA, and Davis TA. Early Characterization of Blast-related Heterotopic Ossification in a Rat Model. Clin Orthop Relat Res. 2015;473(9):2831–9.

45. Kaplan FS, Le Merrer M, Glaser DL, Pignolo RJ, Goldsby RE, Kitterman JA, et al. Fibrodysplasia ossificans progressiva. Best Practice & Research in Clinical Rheumatology. 2008;22(1):191–205.

46. Pignolo RJ, Bedford-Gay C, Liljesthrom M, Durbin-Johnson BP, Shore EM, Rocke DM, et al. The Natural History of Flare-Ups in Fibrodysplasia Ossificans Progressiva (FOP): A Comprehensive Global Assessment. J Bone Miner Res. 2016;31(3):650–6.

47. Leslie AG. The integration of macromolecular diffraction data. Acta crystallographica Section D, Biological crystallography. 2006;62(Pt 1):48–57.

48. Evans P. Scaling and assessment of data quality. Acta crystallographica Section D, Biological crystallography. 2006;62(Pt 1):72–82.

49. McCoy AJ, Grosse-Kunstleve RW, Adams PD, Winn MD, Storoni LC, and Read RJ. Phaser crystallographic software. J Appl Crystallogr. 2007;40(Pt 4):658–74.

50. Emsley P, and Cowtan K. Coot: model-building tools for molecular graphics. Acta Crystallographica Section D-Biological Crystallography. 2004;60:2126–32.

51. Murshudov GN, Vagin AA, and Dodson EJ. Refinement of macromolecular structures by the maximum-likelihood method. Acta crystallographica Section D, Biological crystallography. 1997;53(Pt 3):240–55.

52. Adams PD, Afonine PV, Bunkoczi G, Chen VB, Davis IW, Echols N, et al. PHENIX: a comprehensive Python-based system for macromolecular structure solution. Acta crystallographica Section D, Biological crystallography. 2010;66(Pt 2):213–21.

53. Painter J, and Merritt EA. Optimal description of a protein structure in terms of multiple groups undergoing TLS motion. Acta Crystallographica Section D-Biological Crystallography. 2006;62:439–50.

54. Davis IW, Leaver-Fay A, Chen VB, Block JN, Kapral GJ, Wang X, et al. MolProbity: allatom contacts and structure validation for proteins and nucleic acids. Nucleic acids research. 2007;35(Web Server issue):W375–83.

55. Mullins MC, Hammerschmidt M, Kane DA, Odenthal J, Brand M, van Eeden FJ, et al. Genes establishing dorsoventral pattern formation in the zebrafish embryo: the ventral specifying genes. Development. 1996;123:81–93.

56. Komori T, Yagi H, Nomura S, Yamaguchi A, Sasaki K, Deguchi K, et al. Targeted disruption of Cbfa1 results in a complete lack of bone formation owing to maturational arrest of osteoblasts. Cell. 1997;89(5):755–64.

