## Supplemental Tables and Figures for "Saracatinib is an efficacious clinical candidate for fibrodysplasia ossificans progressiva"

| Inhibitor | ΔTm (°C) ALK2 | ΔTm (°C) ALK5 |
|---|---|---|
| Saracatinib | 13.9 | 10.2 |
| ASP3026 | 10.7 | 3.0 |
| GSK650394 | 10.3 | 10.2 |
| Momelotinib | 9.7 | 5.0 |
| Lestaurtinib | 9.5 | 9.8 |
| AZD1480 | 9.2 | 5.3 |
| (5Z)-7-Oxozeaenol | 9.0 | 5.5 |
| Gandotinib | 8.7 | 7.4 |
| AT9283 | 8.7 | 9.0 |
| LY2874455 | 8.5 | 10.6 |
| Crizotinib | 8.3 | 4.4 |
| Dabrafenib | 7.9 | 12.2 |
| Fedratinib | 7.5 | 2.5 |
| TAK-901 | 7.3 | 8.8 |
| Dasatinib | 6.8 | 6.5 |
| Nintedanib | 6.6 | 7.2 |
| PIK-75 | 6.6 | 8.2 |
| AZD4547 | 5.7 | 1.7 |
| Cabozantinib | 5.4 | 5.3 |
| Baricitinib | 5.1 | 0.5 |
| Danusertib | 5.0 | 2.5 |
| BX-912 | 4.9 | 1.4 |
| PF-431396 | 4.6 | 3.5 |
| Gefitinib | 4.5 | 0.8 |
| Milciclib | 4.5 | 2.5 |
| SP600125 | 4.4 | 1.7 |
| Cediranib | 4.0 | 5.1 |
| Alvocidib | 3.7 | 0.7 |
| Neratinib | 3.6 | 5.1 |
| Tamatinib | 3.6 | 5.0 |
| AP26113 | 3.5 | 3.2 |
| GSK269962 | 3.4 | -1.0 |
| MGCD-265 | 3.4 | 0.6 |
| Ruboxistaurin | 3.4 | 0.1 |
| Crenolanib | 3.4 | 3.2 |
| GSK1838705A | 3.3 | 0.3 |
| AZ23 | 3.3 | 3.3 |
| GSK1904529A | 3.2 | 0.4 |
| SCH772984 | 3.2 | -0.1 |
| AZD7762 | 3.2 | 1.3 |
| Volasertib | 3.1 | 0.2 |
| Orantinib | 3.1 | 0.5 |
| CUDC-101 | 3.0 | 0.1 |
| Dovitinib | 2.9 | -0.1 |
| Voxtalisib | 2.9 | 0.0 |
| Linifanib | 2.7 | 3.3 |
| Sotrastaurin | 2.7 | -0.2 |
| Rigosertib | 2.7 | -0.1 |
| AZD2014 | 2.7 | 0.3 |
| Nilotinib | 2.6 | 1.1 |
| Tideglusib | 2.6 | -0.4 |
| Axitinib | 2.6 | 4.7 |
| INK128 | 2.5 | -0.2 |
| Galunisertib | 2.5 | 11.4 |
| Rebastinib | 2.4 | 0.1 |
| Lenvatinib | 2.3 | 2.5 |
| Pictilisib | 2.3 | 1.1 |
| BMS-599626 | 2.3 | 2.7 |
| Regorafenib | 2.2 | 0.6 |
| Tivozanib | 2.1 | 1.9 |
| Ralimetinib | 2.1 | 5.0 |
| GSK2636771 | 2.0 | 2.7 |
| Ponatinib | 1.8 | 3.1 |
| Dacomitinib | 1.6 | 0.9 |
| Sunitinib | 1.6 | 1.6 |
| PF-670462 | 1.5 | 7.1 |
| BYL719 | 1.5 | 1.0 |
| Ruxolitinib | 1.4 | 1.2 |
| PF-00477736 | 1.4 | 1.4 |
| Vandetanib | 1.3 | 0.7 |
| AZD1152-HQPA | 1.3 | 0.9 |
| Enzastaurin | 1.1 | 1.7 |
| Bosutinib | 1.0 | 1.4 |
| Dinaciclib | 0.9 | 1.8 |
| AZD5363 | 0.8 | 0.2 |
| Alectinib | 0.7 | 2.6 |
| Brivanib | 0.6 | 2.4 |
| BMS-754807 | 0.6 | 1.3 |
| Palbociclib | 0.5 | 0.1 |
| PF-4800567 | 0.5 | 3.5 |
| Motesanib | 0.5 | 0.2 |
| Midostaurin | 0.4 | 1.1 |
| BMS-911543 | 0.4 | 0.3 |
| OSI-027 | 0.3 | 1.9 |
| Bafetinib | 0.3 | 0.4 |
| Buparlisib | 0.3 | 0.6 |
| Afatinib | 0.3 | 0.0 |
| Ibrutinib | 0.2 | 3.1 |
| Dactolisib | 0.2 | 0.7 |
| Pazopanib | 0.1 | 2.1 |
| Amuvatinib | 0.1 | 1.8 |
| AZD8330 | 0.1 | 0.2 |
| RAF265 | 0.0 | 1.7 |
| SNS-032 | 0.0 | 0.0 |
| KX2-391 | 0.0 | 0.5 |
| Ipatasertib | 0.0 | 0.2 |
| Selumetinib | 0.0 | -0.1 |
| Idelalisib | 0.0 | 0.2 |
| PF-04691502 | -0.1 | -0.1 |
| Linsitinib | -0.1 | 0.1 |
| Lapatinib | -0.1 | 0.4 |
| AZ3146 | -0.1 | -0.2 |
| Imatinib | -0.2 | 0.0 |
| ZSTK474 | -0.2 | 0.4 |
| PF-03758309 | -0.2 | 0.0 |
| GSK2126458 | -0.2 | 1.4 |
| EMD1214063 | -0.2 | -0.2 |
| GSK2334470 | -0.3 | -0.3 |
| NVP-BGJ398 | -0.3 | -0.1 |
| PF-04708671 | -0.4 | -0.1 |
| MK-2206 | -0.4 | -0.2 |
| Erlotinib | -0.4 | 0.9 |
| AZD8055 | -0.4 | -0.1 |
| Rabusertib | -0.4 | -0.1 |
| BI 2536 | -0.5 | -0.2 |
| Tandutinib | -0.5 | -0.3 |
| Foretinib | -0.6 | 0.3 |
| Trametinib | -0.6 | -0.1 |
| VX-11E | -0.6 | 0.0 |
| Apitolisib | -0.6 | 0.4 |
| UCN-01 | -0.7 | 0.6 |
| Alisertib | -0.8 | 0.2 |
| Refametinib | -0.8 | -0.3 |
| TGX-221 | -0.8 | 1.3 |
| MK-1775 | -1.1 | 0.5 |
| Pimasertib | -1.3 | -0.4 |
| Tivantinib | -1.4 | 0.2 |
| Vemurafenib | -1.6 | 0.2 |
| Sonolisib | -1.6 | 0.3 |
| Fasudil | -1.6 | -0.1 |
| Tofacitinib | -1.7 | -0.1 |
| Icotinib | -1.7 | 0.4 |
| Pilaralisib | -1.8 | -0.1 |
| Masitinib | -1.8 | 0.0 |
| TAK-733 | -2.0 | -0.3 |
| Doramapimod | -2.0 | -0.3 |
| MEK-162 | -2.0 | -0.4 |
| Vatalanib | -2.1 | 0.0 |
| Seliciclib | -2.1 | -0.3 |
| Apatinib | -2.1 | -0.3 |
| IPI-145 | -2.3 | 0.0 |
| Sorafenib | -2.4 | 0.4 |
| Quizartinib | -2.5 | -0.3 |

**Supplemental Table 1. DSF screening results for wild-type human ALK2 and ALK5 proteins.** Thermal shift values ($\Delta T_m$) are shown from a screen of ~150 clinically relevant small-molecule kinase inhibitors using differential scanning fluorimetry (DSF). As a reference control, the tool compound LDN-193189 gave a Tm shift of 14.5°C.

| | ALK2-saracatinib PDB ID 6ZGC |
|---|---|
| **Data collection** | |
| Space group | I121 |
| Cell dimensions | |
| *a*, *b*, *c* (Å) | 84.6, 101.7, 180.1 |
| α, β, γ (°) | 90.0, 95.3, 90.0 |
| Resolution (Å) | 2.66-44.8 (2.66-2.77) |
| $R_{merge}$ | 0.079 (0.444) |
| $I/\sigma I$ | 7.8 (2.2) |
| Completeness (%) | 96.5 (98.3) |
| Redundancy | 3.3 (3.4) |
| **Refinement** | |
| Resolution (Å) | 2.66-44.8 |
| No. reflections | 41875 (4448) |
| $R_{work/}$ $R_{free}$ | 21.2/26.0 |
| No. atoms | 9252 |
| Protein | 8996 |
| Ligand/ion | 236 |
| Water | 20 |
| B-factors | 70.2 |
| Protein | 70.3 |
| Ligand/ion | 67.1 |
| Water | 59.9 |
| R.m.s deviations | |
| Bond lengths (Å) | 0.007 |
| Bond angles (º) | 0.844 |
| Ramachandran statistics | |
| Preferred (%) | 98 |
| Allowed (%) | 99.8 |

**Supplemental Table 2** Data collection and refinement statistics (Molecular replacement)
Data from a single crystal.
*Highest resolution shell is shown in parenthesis.

| | % inhib | |
|---|---|---|
| Kinase | 100 nM | 1 μM |
| ABL1 | 93.5 | 99.9 |
| AKT1 | 0.4 | 2.7 |
| AKT2 | 0.2 | 1.6 |
| AKT3 | 1.9 | 2.8 |
| ALK | 12.7 | 55.6 |
| ALK2 | 95.8 | 98.1 |
| ALK4 | -8.4 | 63.4 |
| AMP-α1β1γ1 | 2.1 | 5.6 |
| AMP-α2β1γ1 | -0.8 | 2.5 |
| ARG (ABL2) | 82.5 | 97.5 |
| ARK5 | -0.6 | 10.3 |
| AURORA-A | 0.3 | 3.8 |
| AURORA-B | 1.8 | 33.6 |
| AURORA-C | 5.0 | 6.8 |
| AXL | 5.6 | 30.4 |
| BLK | 81.7 | 98.7 |
| BMX | 9.7 | 43.1 |
| BRAF | -0.3 | -1.9 |
| BRK | 75.0 | 97.1 |
| BRSK1 | 4.4 | -7.5 |
| BRSK2 | -0.9 | 1.8 |
| BTK | 17.0 | 57.3 |
| CAMK1α | 1.6 | 4.0 |
| CAMK1δ | -1.2 | 1.3 |
| CAMK2α | 2.7 | -3.1 |
| CAMK2β | 0.4 | -0.4 |
| CAMK2δ | -1.4 | -2.2 |
| CAMK2γ | 3.8 | 0.7 |
| CAMK4 | 2.8 | 3.4 |
| CDK1/CycB | -2.5 | 2.9 |
| CDK2/CycA | -0.1 | 3.9 |
| CDK2/CycE | 1.2 | 4.2 |
| CDK3/CycE | 2.6 | 6.9 |
| CDK4/CycD | -0.3 | 7.7 |
| CDK5 | 0.9 | 2.9 |
| CDK5/p35 | 7.5 | 9.5 |
| CDK6/CycD3 | 1.2 | 6.5 |
| CDK7 | -1.4 | 4.9 |
| CDK9-Cyc1 | 3.4 | 3.0 |
| CHEK1 | 2.7 | 9.7 |
| CHEK2 | 2.7 | 9.8 |
| CK1α | -0.2 | 9.3 |
| CK1-ε | -0.2 | 15.0 |
| CK1-γ1 | -2.6 | -0.1 |
| CK1-γ2 | -4.7 | -0.2 |
| CK1-γ3 | -3.7 | 1.1 |
| CLK1 | 4.1 | 0.8 |
| CLK2 | 1.8 | 4.7 |
| CLK3 | 6.2 | 6.4 |
| CLK4 | 1.8 | 21.9 |
| CRAF | -8.5 | 2.8 |
| CSK | 21.0 | 70.0 |
| DAPK1 | 0.9 | 1.9 |
| DAPK3 | 0.7 | 0.4 |
| DCAMKL2 | -2.8 | -0.5 |
| DDR1 | 89.6 | 98.3 |
| DDR2 | 71.8 | 96.2 |
| DYRK1A | -0.5 | 2.9 |
| DYRK1B | 1.0 | 1.2 |
| DYRK2 | -0.3 | 0.8 |
| DYRK3 | -1.6 | 1.2 |
| DYRK4 | -2.8 | -0.1 |
| EGFR | 51.8 | 88.9 |

| | % inhib | |
|---|---|---|
| Kinase | 100 nM | 1 μM |
| EPH A1 | 86.5 | 98.1 |
| EPH A2 | 50.4 | 92.6 |
| EPH A3 | 13.9 | 65.3 |
| EPH A4 | 57.0 | 97.2 |
| EPH A5 | 40.9 | 88.1 |
| EPH A8 | 47.9 | 90.3 |
| EPH B1 | 61.3 | 94.4 |
| EPH B2 | 62.4 | 95.4 |
| EPH B3 | -3.8 | 0.0 |
| EPH B4 | 46.1 | 90.5 |
| ERB-B2 | 18.0 | 63.3 |
| ERBB4 | 62.3 | 95.3 |
| FAK | -1.9 | 2.3 |
| FER | -0.2 | 11.4 |
| FES | -1.2 | 2.6 |
| FGFR1 | 3.5 | 13.5 |
| FGFR2 | 0.3 | 19.9 |
| FGFR3 | -3.7 | 3.1 |
| FGFR4 | -1.9 | 2.5 |
| FGR | 67.0 | 95.3 |
| FLT 1 | -9.8 | 18.1 |
| FLT 3 | 12.0 | 56.9 |
| FLT 4 | -0.6 | 5.4 |
| FMS | 53.5 | 90.0 |
| FRAP1 | 6.4 | 7.2 |
| FYN | 80.2 | 100.1 |
| GRK6 | -3.2 | -0.7 |
| GRK7 | -1.8 | 8.0 |
| GSK3α | 0.1 | 4.0 |
| GSK3β | 0.2 | 1.8 |
| HASPIN | 0.9 | -0.8 |
| HCK | 73.4 | 83.7 |
| HIPK1 | -1.8 | 0.7 |
| HIPK2 | 0.6 | 1.9 |
| HIPK3 | -3.5 | -1.8 |
| HIPK4 | 0.0 | 15.7 |
| IGF1R | -2.1 | 4.2 |
| IKK α | 0.8 | 8.7 |
| IKK β | -0.9 | -3.6 |
| IKK ε | -1.6 | 7.3 |
| INSR | -0.8 | 3.8 |
| IRAK1 | 13.6 | 37.8 |
| IRAK4 | 0.0 | 20.5 |
| IRR | -1.5 | -2.8 |
| ITK | 4.0 | 15.6 |
| JAK1 | 2.0 | 0.8 |
| JAK2 | -5.1 | 6.2 |
| JAK3 | -0.1 | -0.6 |
| JNK1 | 0.1 | 3.4 |
| JNK2 | -1.2 | 3.1 |
| JNK3 | -2.7 | 0.2 |
| KDR | 3.3 | 9.3 |
| KIT | 52.9 | 89.1 |
| LATS1 | -0.5 | 6.1 |
| LATS2 | 1.5 | 13.1 |
| LCK | 97.6 | 100.4 |
| LOK | 33.8 | 84.2 |
| LRRK2 | 3.1 | 35.0 |
| LTK | 17.1 | 73.6 |
| LYNA | 83.5 | 98.9 |
| LYNB | 81.4 | 98.2 |
| MAP4K2 | 3.2 | 14.0 |
| MAP4K4 | 32.5 | 84.7 |

| | % inhib | |
|---|---|---|
| Kinase | 100 nM | 1 μM |
| MAP4K5 | 41.5 | 102.1 |
| MAPK1 | -0.4 | -1.2 |
| MAPK3 | -1.7 | 0.6 |
| MAPKAPK2 | -1.5 | 3.6 |
| MAPKAPK3 | 1.8 | 22.5 |
| MARK1 | -2.2 | 0.4 |
| MARK3 | -1.5 | 1.4 |
| MARK4 | -0.8 | 2.4 |
| MEK1 | -1.1 | -0.6 |
| MEK2 | -1.1 | 1.2 |
| MELK | -0.2 | 5.1 |
| MER | 4.0 | 19.7 |
| MET | 3.4 | 4.9 |
| MINK | 19.4 | 73.3 |
| MKNK1 | -1.9 | 5.7 |
| MRCK α | 1.9 | 2.8 |
| MRCK β | -0.2 | 0.6 |
| MSK1 | -3.2 | 0.4 |
| MSK2 | -2.6 | 1.8 |
| MSSK1 | 3.3 | 2.7 |
| MST1 | 1.3 | 7.5 |
| MST2 | 1.7 | 3.5 |
| MST3 | 1.0 | 16.6 |
| MST4 | 8.3 | 12.5 |
| MUSK | 2.5 | 17.5 |
| NDR1 | -2.3 | 0.9 |
| NDR2 | -0.8 | 3.1 |
| NEK1 | -1.9 | 4.2 |
| NEK2 | 0.2 | 1.5 |
| NEK6 | -4.1 | -1.1 |
| NEK7 | -0.5 | 1.8 |
| NEK9 | -0.4 | 0.9 |
| P38α | 10.5 | 54.9 |
| P38β | -1.3 | 0.1 |
| P38γ | 0.0 | 2.7 |
| P38δ | -1.1 | 0.4 |
| P70S6K1 | 0.2 | 6.5 |
| P70S6KB1 | 1.1 | 1.0 |
| PAK1 | -0.5 | 1.9 |
| PAK2 | 0.3 | 3.2 |
| PAK3 | -1.5 | -0.3 |
| PAK4 | 1.1 | 7.7 |
| PAK5 | 1.7 | 5.3 |
| PAK6 | 1.3 | 2.1 |
| PAR-1Bα | 2.1 | 5.9 |
| PASK | 0.6 | -3.5 |
| PDGFRα | 31.8 | 85.6 |
| PDGFRβ | 53.7 | 91.8 |
| PDK1 | -1.8 | 1.8 |
| PERK | 10.0 | 4.0 |
| PHKγ1 | 1.0 | 9.2 |
| PHKγ2 | -2.6 | 5.3 |
| PI3-K-α | -10.3 | 4.9 |
| PI4-K-β | 1.5 | 2.6 |
| PIM 1 | -0.7 | 3.2 |
| PIM 2 | -0.5 | -1.4 |
| PIM3 | -0.3 | -0.2 |
| PKA | 1.0 | 6.2 |
| PKACB | -2.0 | 0.6 |
| PKC-α | 2.8 | 7.2 |
| PKC-β1 | 2.2 | 5.4 |
| PKC-β2 | 1.3 | -0.3 |
| PKC-ε | 2.2 | 13.0 |

| | % inhib | |
|---|---|---|
| Kinase | 100 nM | 1 μM |
| PKC-η | -2.0 | 0.7 |
| PKC-γ | -2.1 | 2.4 |
| PKC-ι | -8.3 | 2.8 |
| PKC-θ | -0.6 | 4.8 |
| PKC-η | 2.5 | 13.7 |
| PKN1 | 1.0 | 0.6 |
| PKN2 | -2.9 | -1.9 |
| PLK1 | -5.7 | -5.1 |
| PLK3 | -12.2 | -7.0 |
| PLK4 | 2.2 | -7.3 |
| PRAK | 0.9 | 4.8 |
| PRKACA | -1.8 | 1.4 |
| PRKD1 | -1.9 | 4.3 |
| PRKD2 | -0.6 | 3.7 |
| PRKD3 | -2.6 | 6.5 |
| PRKG1 | 3.1 | 6.9 |
| PRKG2 | -4.8 | 8.5 |
| PRKX | -1.1 | 2.1 |
| PTK5 | 23.3 | 75.3 |
| PYK2 | 11.8 | 29.4 |
| RET | 6.6 | 34.2 |
| RIPK2 | 93.8 | 95.8 |
| ROCK1 | 1.8 | 6.9 |
| ROCK2 | -0.1 | 9.1 |
| RON | 0.4 | 11.8 |
| ROS | 1.2 | 22.5 |
| RSK1 | 1.4 | 6.3 |
| RSK2 | 2.0 | 6.1 |
| RSK3 | -4.6 | 1.1 |
| RSK4 | 0.6 | 2.2 |
| SGK1 | -0.2 | 1.0 |
| SGK2 | 1.8 | 3.3 |
| SGK3 | -0.2 | 0.3 |
| SLK | 31.7 | 85.6 |
| SIK | 10.8 | 59.9 |
| SIK2 | 46.2 | 86.7 |
| SPHK1 | -0.7 | -4.7 |
| SPHK2 | -10.2 | -11.7 |
| SRC | 94.1 | 100.8 |
| SRMS | 3.2 | 12.7 |
| SRPK1 | -1.4 | -0.6 |
| SRPK2 | 6.1 | 5.0 |
| STK16 | -10.0 | 1.7 |
| SYK | 0.6 | -4.5 |
| TAK1-TAB1 | -1.0 | 2.9 |
| TAOK2 | -1.0 | 3.4 |
| TAOK3 | 5.9 | 5.6 |
| TBK1 | -0.4 | 4.6 |
| TEC | 8.4 | 35.4 |
| TIE2 | 1.9 | -0.8 |
| TNIK | 50.4 | 96.7 |
| TNK2 | 35.0 | 87.0 |
| TRKA | 6.0 | 25.5 |
| TRKB | 3.8 | 17.4 |
| TRKC | 4.6 | 17.9 |
| TSSK1 | 2.6 | 10.5 |
| TSSK2 | 2.8 | 4.0 |
| TTK | 1.4 | 2.7 |
| TXK | 41.8 | 87.0 |
| TYK2 | 3.3 | 0.4 |
| TYRO3 | 11.8 | 57.8 |
| YES | 92.8 | 99.6 |
| ZAP70 | 0.8 | 5.4 |

**Supplemental Table 3**. **Saracatinib selectivity profiling.** % inhibition values are shown at 100 nM or 1 μM saracatinib using Caliper screening (Nanosyn).

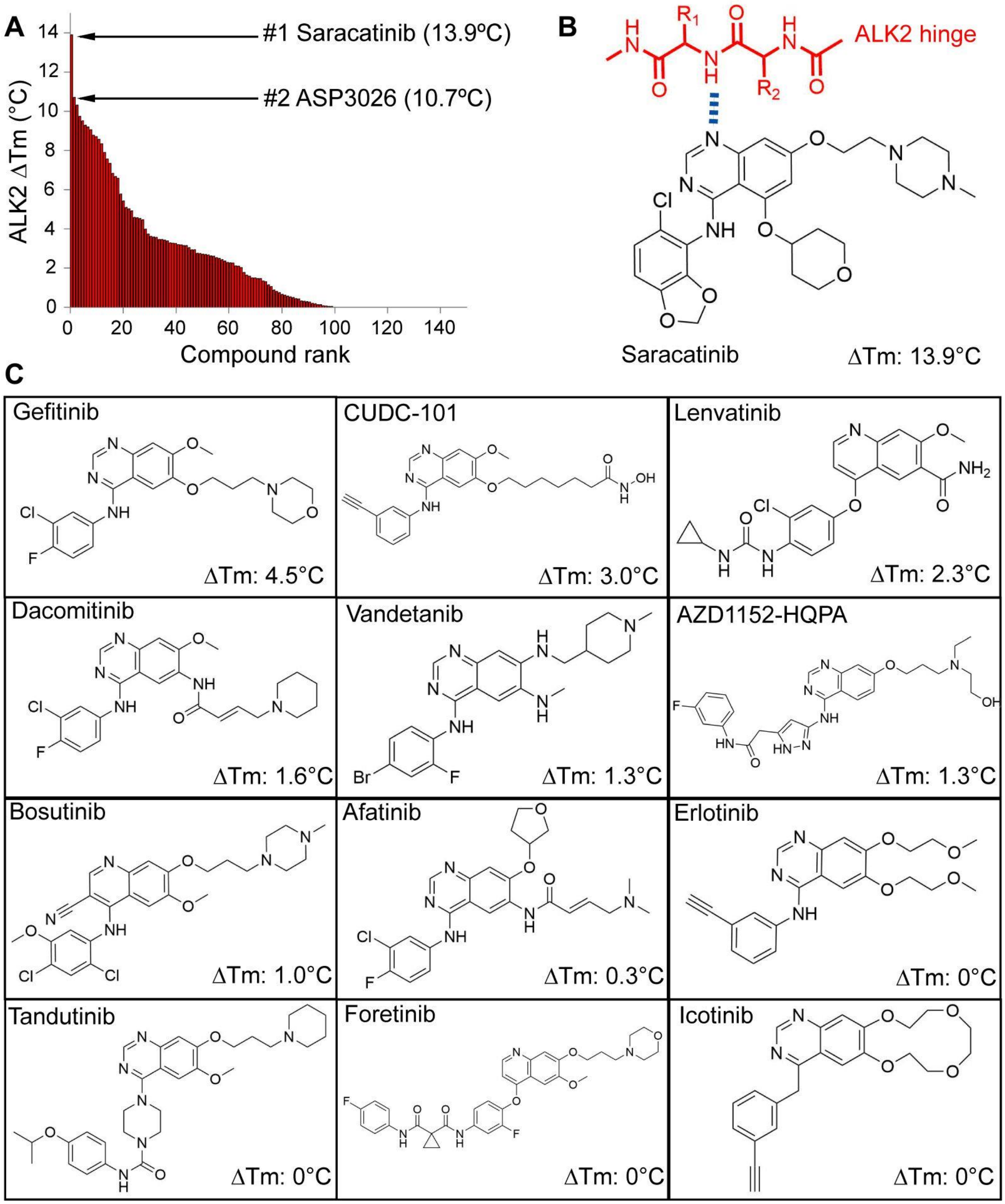

**Supplemental Figure 1**. **Comparison of ALK2 binding to different quinazoline and quinoline scaffolds.** (**A**) Compounds were ranked for their binding to ALK2 according to their thermal shift values ($\Delta T_m$). (**B**) Saracatinib chemical scaffold highlighting the position of the quinazoline hydrogen bond to the kinase hinge region. (**C**) Chemical scaffolds and ALK2 $\Delta T_m$ values obtained with other quinazoline or quinoline-containing compounds in the screening panel.

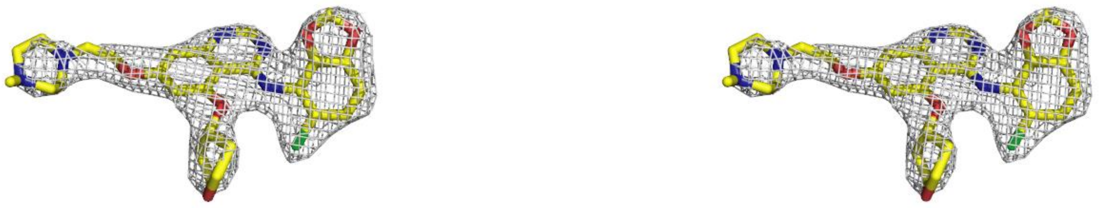

**Supplemental Figure 2. Electron density for saracatinib.** Stereo view of the electron density (Fo-Fc) map of saracatinib at σ=2.0.

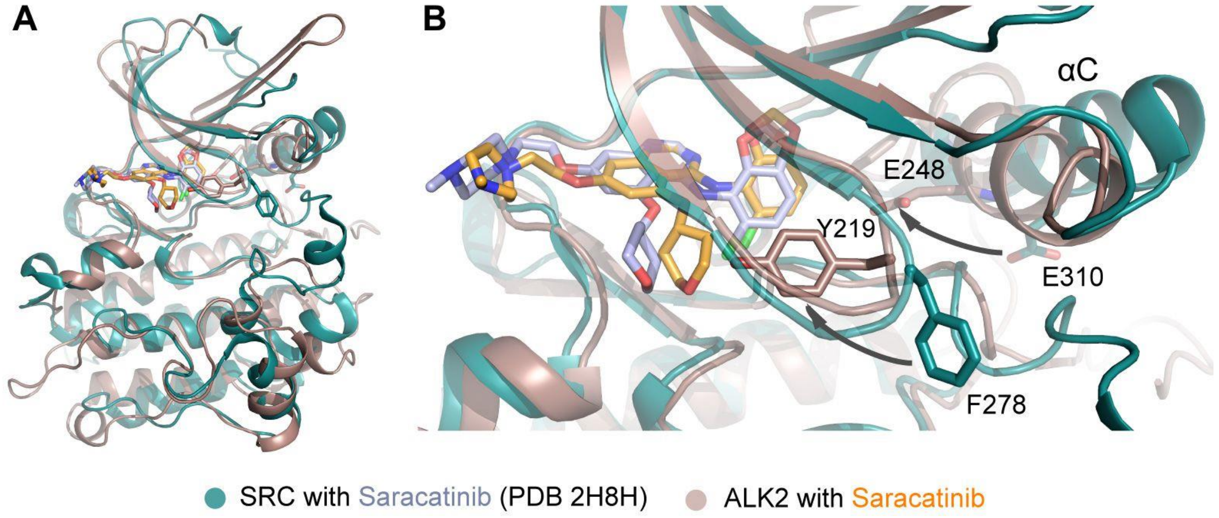

**Supplemental Figure 3. Saracatinib interactions with SRC and ALK2.** (**a**) Superposition of the saracatinib complexes with SRC (PDB 2H8H) and ALK2. (**b**) The inactive conformations of the bound SRC and ALK2 kinases are strikingly different. The saracatinib complex with SRC shows an αC-out conformation such that SRC Glu310 is rotated away from the ATP-binding pocket. In contrast, ALK2 shows an inactive αC-in conformation in which the equivalent αC residue Glu248 is swung inwards. ALK2 also shows an inactive conformation of the glycine-rich loop which positions Tyr219 inside the ATP pocket. Arrows highlight the different positions of selected side chains.

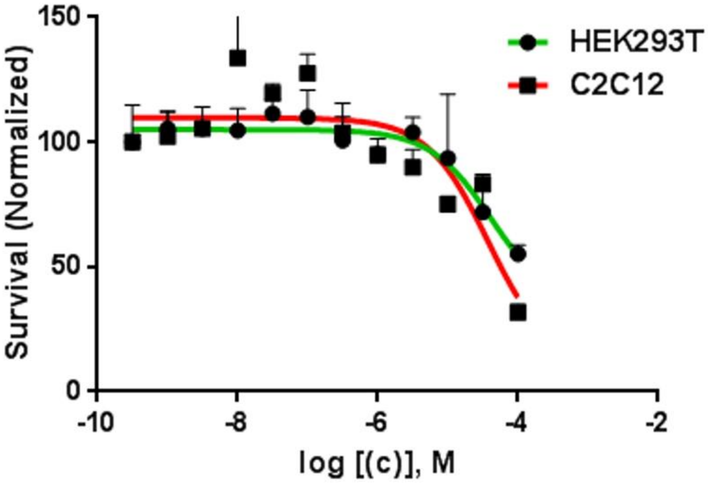

| | $ED_{50}$ |
|---|---|
| HEK293T | 4.19E-05 |
| C2C12 | 3.75E-05 |

**Supplemental Figure 4. Tolerability of saracatinib**. Tolerability of saracatinib in indicated cell lines was determined using the MTT cell viability assay. Data were plotted as mean + S.E.M. and the average $ED_{50}$ determined (n=3 replicates).

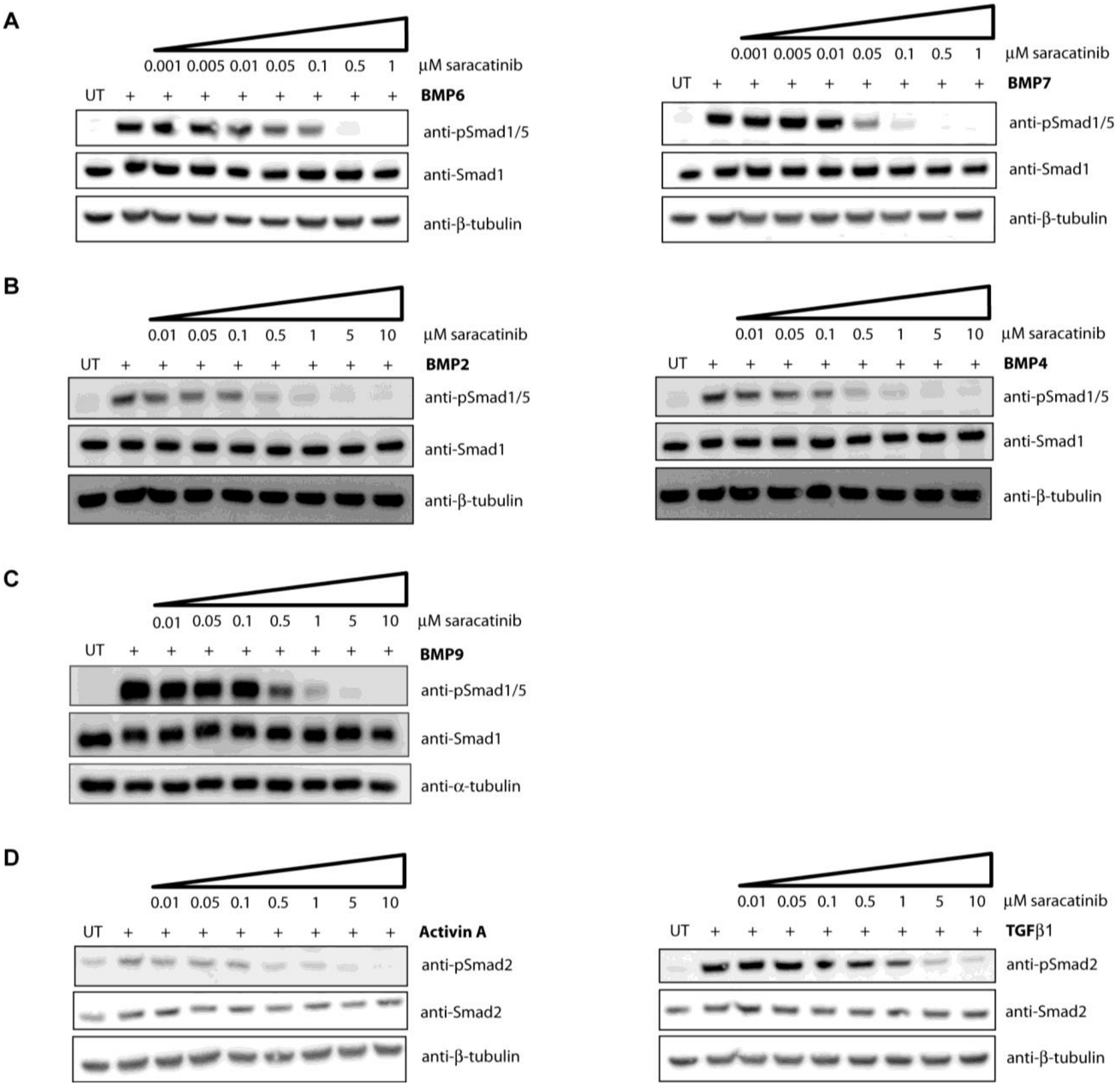

**Supplemental Figure 5. Saracatinib inhibition of Smad phosphorylation.** (**A**) Western blot analysis showing potent saracatinib inhibition of Smad1/5 phosphorylation following BMP6 or BMP7 stimulation in C2C12 cells. (**B**) Saracatinib inhibition of Smad1/5 phosphorylation following BMP2 or BMP4 stimulation in C2C12 cells, or (**C**) BMP9 stimulation in bovine aortic endothelial cells (BAEC). (**D**) Saracatinib inhibition of Smad2 phosphorylation following TGFβ1 or activin A stimulation in C2C12 cells.

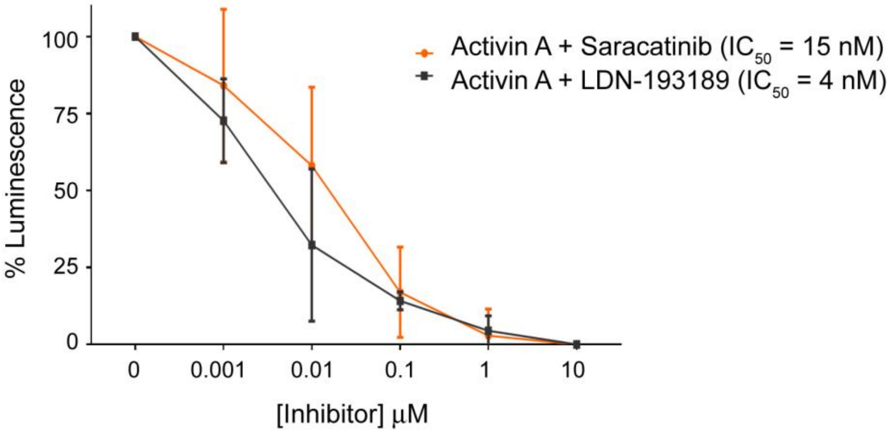

**Supplemental Figure 6. In-cell immunofluorescent assay of phospho-SMAD1/5 levels.** An in-cell chemiluminescent assay confirmed that both Saracatinib and LDN-193189 inhibit the neofunction of ALK2$^{R206H}$ in FOP patient-derived dermatofibroblasts by diminishing activin A mediated pSMAD1/5 signaling. Cells were pre-incubated with increasing molar concentrations of saracatinib or LDN-193189 for 15 minutes and then treated with 250 ng/mL activin A for a further 45 minutes. Following fixation, cells were incubated with primary antibody (Phospho-Smad1/5 (Ser463/465) Rabbit mAb #9516; Cell Signaling Technologies, MA) and secondary antibody (Anti-rabbit IgG, HRP-linked antibody; Cell Signaling Technologies, MA). Finally, the cells were developed with BioFX Chemiluminescent Ultra Sensitive HRP Microwell Substrate solution (Surmodics, MN) and read on a Spectra Max L plate reader (Molecular Devices, CA). Data shown are plotted as mean ± S.D. (n=2 replicates).

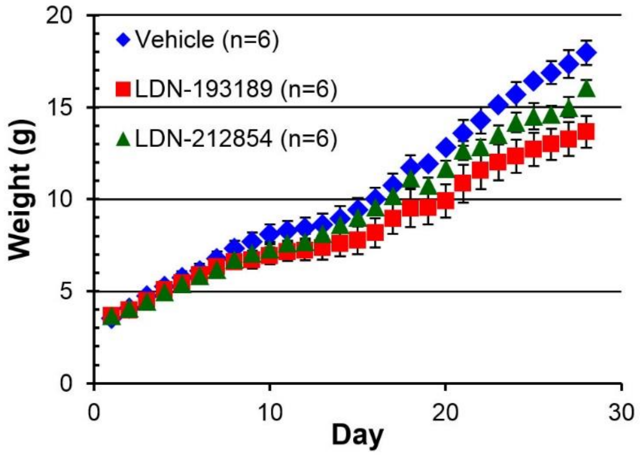

**Supplemental Figure 7. In vivo tolerability of LDN-193189 and LDN-212854 in a mouse model of fibrodysplasia ossificans progressiva (FOP).** Mice expressing an inducible constitutively-active *ACVR1*$^{Q207D}$ (CAG-Z-EGFP-caALK2) transgene were treated with vehicle, or pyrazolo-pyrimidine derivatives LDN-193189 or LDN-212854 (6 mg/kg intraperitoneally twice daily). Progressive heterotopic ossification and passive range-of-motion loss were observed in vehicle-treated mice starting on day 10 after injection of Ad.Cre, whereas mobility was almost entirely preserved in mice treated with LDN-193189 and LDN-212854 (not shown). In contrast to treatment with saracatinib, treatment with LDN-193189 or LDN-212854 was associated with weight loss of between 10% - 25% relative to weights of vehicle-treated controls.

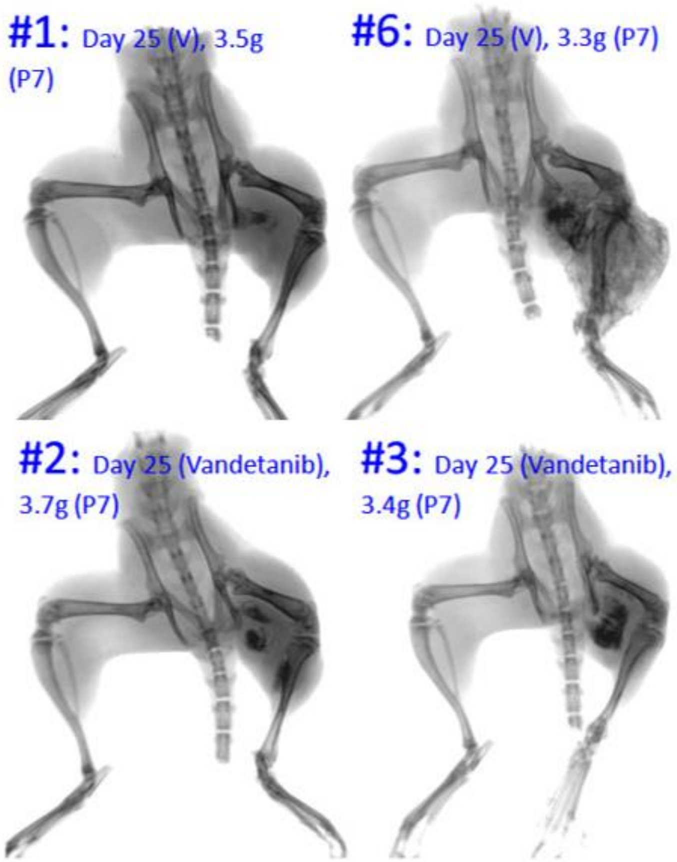

**Supplemental Figure 8. Lack of in vivo efficacy of vandetanib (AZD6474) in a mouse model of heterotopic ossification (HO).** Mice expressing an inducible constitutively-active *ACVR1*$^{Q207D}$ (CAG-Z-EGFP-caALK2) transgene were treated with vehicle, or vandetanib (25 mg/kg intraperitoneally twice daily). Heterotopic ossification was assessed radiographically following injection of Ad.Cre. All animals treated with vehicle developed progressive heterotopic ossification following challenge with Ad.Cre, as in previous studies. There was no evidence of improvement in radiographic HO or range-of motion in mice treated with vandetanib as compared to those treated with vehicle (n=3 each group). Representative mice are shown; vehicle-treated mice on the top panel are indicated by (v).
